# Targeting NF-κB epigenetic activation and DNA repair deficiency in G34-mutant pediatric diffuse hemispheric glioma with nanoparticles combining PARP inhibition and immune stimulation mediated by CpG dinucleotides

**DOI:** 10.64898/2026.03.09.710503

**Authors:** Santiago Haase, Kaushik Banerjee, Anzar Abdul Mujeeb, Troy Halseth, Lisha Liu, Minzhi Yu, Sushmitha Sriramulu, Maya Sheth, Sadhakshi Raghuram, Pedro R. Lowenstein, Anna Schwendeman, Maria G. Castro

## Abstract

Diffuse hemispheric gliomas (DHGs) are highly aggressive and infiltrative CNS tumors that are refringent to treatment, and with a 5-year overall survival of around 20%. A fraction of DHGs is driven by mutations in the histones H3.1 and H3.3. In this study, we demonstrate that the expression of histone H3.3 glycine 34 to arginine mutations (H3.3-G34R) result in the epigenetic and transcriptional activation of the NF-κB signaling pathway in DHG. To target this vulnerability, we designed high density lipoprotein (HDL) nanoparticles loaded with unmethylated CpG dinucleotides, which mimic the immune stimulatory activity of bacterial DNA. CpG are recognized by Toll-like receptor 9 (TLR9), activating the NF-κB signaling. The CpG-mediated NF-κB activation results in the release of immuno-stimulating cytokines that promote an antitumoral response. As we previously established that G34-mutant DHGs are characterized by DNA repair impairment, we combined CpG dinucleotides with a PARP (poly (ADP-ribose) polymerase) inhibitor, olaparib, in the HDL nanoparticles.

## Introduction

Central nervous system (CNS) are the primary cause of cancer-related fatalities in individuals between the ages of 0 and 14 ^[1]^. These tumors have an incidence of 5.74 per 100,000 population, and a mortality rate of 4.42 per 100,000 population ^[1]^. Diffuse hemispheric gliomas (DHGs) are amongst the most frequent and aggressive pediatric CNS tumors. DHG encompass anaplastic astrocytoma (CNS WHO grade III), glioblastoma multiforme (GBM; CNS WHO grade IV), and diffuse midline gliomas (CNS WHO grade IV), and are characterized for being highly infiltrative, invasive, and aggressive ^[2–4]^. Regrettably, they remain resistant to treatments, with an overall 5-year survival rate of under 20% ^[3,5]^. Even with such alarming statistics, the last ten years have seen substantial progress in understanding the molecular characteristics of DHGs ^[6–8]^. A significant advancement in pediatric glioma biology was the identification of mutations in the histone gene H3. These mutations play an active role in glioma development, correlating with distinct genomic alterations, epigenetic landscapes, CNS locations, and clinical features ^[6–11]^.

Our study centers on the subtype of DHG driven by a mutation in the histone H3.3 gene H3F3A, which alters the glycine at position 34 to an arginine (H3.3-G34R), or, less frequently, to a valine ^[6,7]^. This mutation is observed in 9-15% of DHGs, is limited to the cerebral hemispheres, and predominantly occurs in the adolescent demographic (median 15.0 years) ^[6–8,12,13]^. G34-mutant DHGs are hallmarked by the presence of loss of function mutations in ATRX/DAXX and TP53 genes, and a DNA hypomethylated phenotype ^[6–8]^.

These tumors pose a formidable prognosis, with patients displaying a median survival of 18.0 months and a 2-year overall survival rate of 27.3% ^[6]^.

The mechanisms underlying disease progression in the H3.3-G34R/V subgroup of diffuse hemispheric gliomas (DHGs) are not completely understood ^[8,14,15]^. Presently, the standard care for non-midline DHGs, irrespective of their H3 status, includes maximum surgical removal followed by targeted radiation. The potential benefit of adding chemotherapy to radiation therapy for G34-mutant DHGs is still a topic of contention, largely because earlier cooperative group studies did not consistently test and stratify for G34 mutations ^[16]^.

The introduction of effective immunotherapies for DHG is in its early stages. Recent studies in diffuse hemispheric gliomas (DHG) have highlighted a complex and dynamic interaction between the tumor cells and the immune system ^[17]^. DHGs are heterogeneous tumors with a significant portion of the tumor mass composed of non-malignant cells, such as macrophages, microglia, and dendritic cells, as well as myeloid-derived suppressor cells (MDSCs). These have been identified as major contributors to immunosuppression within the tumor microenvironment ^[18,19]^. Consequently, the development of immunotherapies has been hindered by several factors, including the anti-inflammatory tumor microenvironment often observed in these tumors, which is influenced by numerous factors, including the specific tumor subtype, the genetic aberrations present within the tumor, and the host immune system. We recently reported that G34 mutations in DHG lead to epigenetic activation of immune-related genes in the tumor cells which result in a functional antitumoral immune activation of the TME, making these tumors more immunogenic ^[20]^. These results align with recent single-cell transcriptomic studies, where G34-mutant DHGs were shown to feature a relatively small population of immune cells, unlike adult gliomas that are characterized by a large count of immune cells exhibiting an immunosuppressive phenotype ^[21]^. Additionally, a recent study documented the impact of germinal G34 mutations on mouse phenotypes ^[22]^. The authors found that the G34-mutant histone leads to a reduction in H3K36me2, which subsequently results in the hindered recruitment of DNMT3A, a DNA methyltransferase, *in cis*. This, in turn, causes hypomethylation in the DNA, notably impacting the regulatory regions of immune-related genes. This hypomethylation triggers an increase in the expression of these genes, leading to the activation of proinflammatory signals in the brains of G34-mutant mice ^[22]^. These findings correlated with the accumulation of microglia/astrocytes and neuronal depletion in G34-mutant mice. All these results indicate that the G34-mutations correlate with a more immunoreactive TME phenotype.

Our previous results also demonstrate that G34-mutations cause impaired DNA repair to DHG, and that due to this, G34-mutant DHG are more susceptible to DNA damaging therapies (including IR) and to DNA damage response (DDR) inhibitors, such as PARP inhibitors ^[16]^. The DNA repair impaired phenotype also causes the release of DNA to the cytoplasm and the activation of the cGAS-STING pathway in G34-mutant DHG, which leads to the release of immuno-stimulatory cytokines ^[16]^.

In this study, we aimed at developing a tailored therapy to target enhanced immune responsiveness and DNA repair impairment in G34-mutant DHG. We identify here that G34-mutations also correlate with an exacerbated upregulation of the NF-κB signaling pathway. The activation of the NF-κB (Nuclear Factor kappa-light-chain-enhancer of activated B cells) pathway plays a pivotal role in the immune response to cancer. This complex is crucial to produce various pro-inflammatory cytokines, chemokines, and cell adhesion molecules that regulate immune cell trafficking and interactions. Consequently, its activation is integral to immune stimulation, aiding in the recruitment, activation, and differentiation of immune cells such as T-cells, B-cells, and natural killer (NK) cells that are essential for an effective anti-tumor response. To exploit the enhanced activation of the NF-κB pathway in G34-mutant DHG, we developed sHDL nanodiscs loaded with unmethylated CpG dinucleotides, and olaparib, a PARP inhibitor. CpG oligodeoxynucleotide is a TRL9 ligand that mimics bacterial DNA and has been shown to trigger immune rejection and induce long-term immunity against gliomas ^[23]^.

## Results

### Development and characterization of sHDL Nanodiscs Loaded with CpG deoxynucleotides and olaparib

To target the activation of the NF-κB pathway in G34-mutant DHG, we developed HDL-mimicking nanodiscs loaded with CpG deoxynucleotides and the PARP inhibitor olaparib. The efficacy of sHDL nanoparticles to deliver chemotherapeutics agents to glioma has been reported by our group ^[23]^. Dynamic light scattering (DLS) and gel permeation chromatography (GPC) were used to examine particle size, homogeneity, and purity of nanodiscs. ApoA-I mimetic peptide (22A), phospholipids (1,2-dimyristoyl-sn-glycero-3-phosphocholine (DMPC) and 1-palmitoyl-2-oleoylglycero-3-phosphocholine (POPC)), and therapeutic compounds CpG and olaparib were combined at a 1:1:1:0.06 weight ratio in an organic solvent, lyophilized, and hydrated with aqueous buffer (Figure 1A-B). The mixture was first heated and then cooled to facilitate particle assembly. Formation of homogeneous nanodiscs with average size of 10–12 nm and was observed (Figure 1C-D) by size exclusion chromatography and transmission electron microscopy (Figure 1E). The nanodisc size determination by DLS correlated with the GPC measurement, and as the size of nanodisc increased, the retention time of GPC peak decreased. Olaparib and CpG were successfully incorporated in sHDL at ∼2% (w/w) loading.

**Figure 1.**
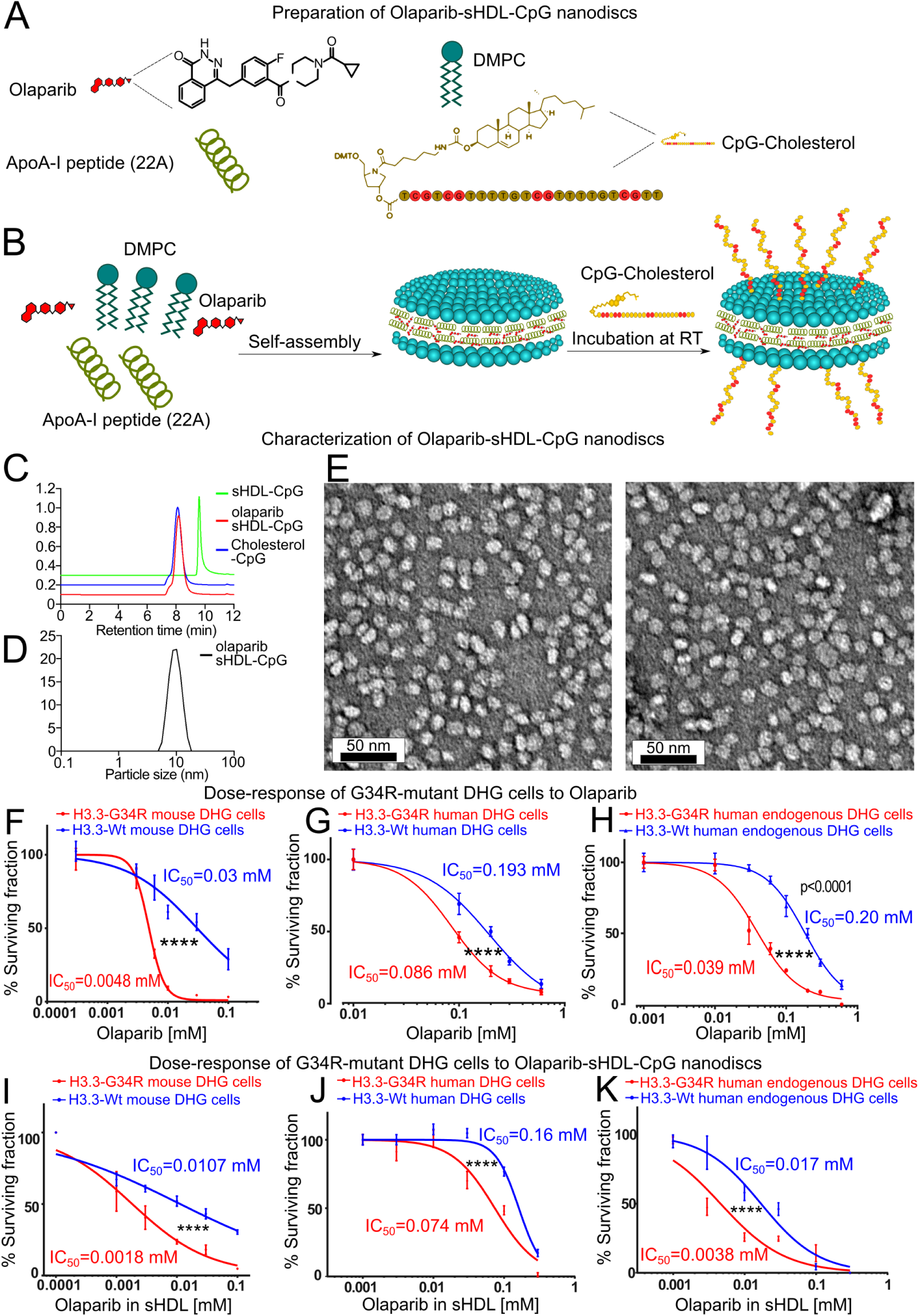
Characterization of Olaparib-sHDL-CpG nanodiscs. A-B) Scheme of Olaparib-sHDL-CpG nanoparticles generation. Olaparib-sHDL-CpG was formulated by first incubating Olaparib with Sphingomyelin and ApoA-I peptide (22A) to form the sHDL with Olaparib. Then the CpG-Cholesterol was added to the already made sHDL discs to form the final disc, Olaparib-sHDL-CpG. C) Characterization of Olaparib-sHDL-CpG nanodiscs. C) Particle size distribution of sHDL-CpG, Olaparib sHDL-CpG, and Cholesterol-CpG determined by gel permeation chromatography (GPC). D) Particle size distribution of Olaparib sHDL-CpG determined by dynamic light scattering (DLS). sHDL concentration of 1 mg/mL. (E) Images demonstrating particle size distribution and morphology of sHDL-CpG and Olaparib-sHDL-CpG taken by transmission electron microscopy (scale bar = 50 µm). G-H) Dose response curves of G34-mutant versus histone wild-type mouse, human and endogenous human diffuse hemispheric glioma cells in response to Olaparib. I-K) Dose response curves of G34-mutant versus histone wild-type mouse, human and endogenous human diffuse hemispheric glioma cells in response to Olaparib-sHDL-CpG nanodiscs.

To assess the functionality of the nanoparticles, we evaluated the survival of H3.3-G34R versus H3.3-Wt DHG cells *in vitro* upon treatment with olaparib-sHDL-CpG nanoparticles in dose-response experiments. Our results demonstrate that mouse DHG cells, human genetically engineered DHG cells and endogenously mutant patient-derived DHG cells expressing the G34 mutation are more susceptible to the nanoparticles than the histone wild-type counterparts (Figure 1F-K). We reported that G34-mutant DHG are characterized by DNA repair impairment ^[16,24]^, which explains the increased susceptibility to the PARP inhibitor. The IC50 concentrations of the nanoparticles are similar to the IC50 concentrations obtained with the non-formulated (free) olaparib (Figure 1F-K), indicating that the incorporation of olaparib into the nanoparticles did not affect the bioactivity of the PARP inhibitor.

### The SR-BI receptor mediates olaparib-sHDL-CpG delivery to glioma cells

Scavenger Receptor Class B Type I (SR-BI) mediates the uptake of HDL into cells (Figure 2A) ^[25]^. The analysis of transcriptional expression of SR-BI (*SCARB1* gene) in GBM versus normal brain tissue from public data available at Gliovis ^[26]^ shows that SR-BI expression is significantly higher in glioma cells than in normal brain cells (Figure 2B). We analyzed SR-BI protein expression by immunofluorescence in H3.3-Wild type and H3.3-G34R mutant mouse and human pediatric DHG cells and astrocytes and show that SR-BI is highly expressed in DHG cells and absent in the astrocyte cells (Figure 2C). We confirmed the immunofluorescence results by western blotting (Figure 2D). Finally, to confirm that SRBI is expressed in our DHG in vivo models, we performed SR-BI immunohistochemistry on fixed brain tissue from DHG tumor-bearing mice to compare SR-BI expression in the tumor versus the tumor-free contralateral hemisphere (Figure 2E-H), confirming SR-BI presence in the tumor and absence in the tumor-free brain. Altogether, this data indicates that the expression of the HDL receptor, SR-BI, is highly enriched in tumor cells.

**Figure 2.**
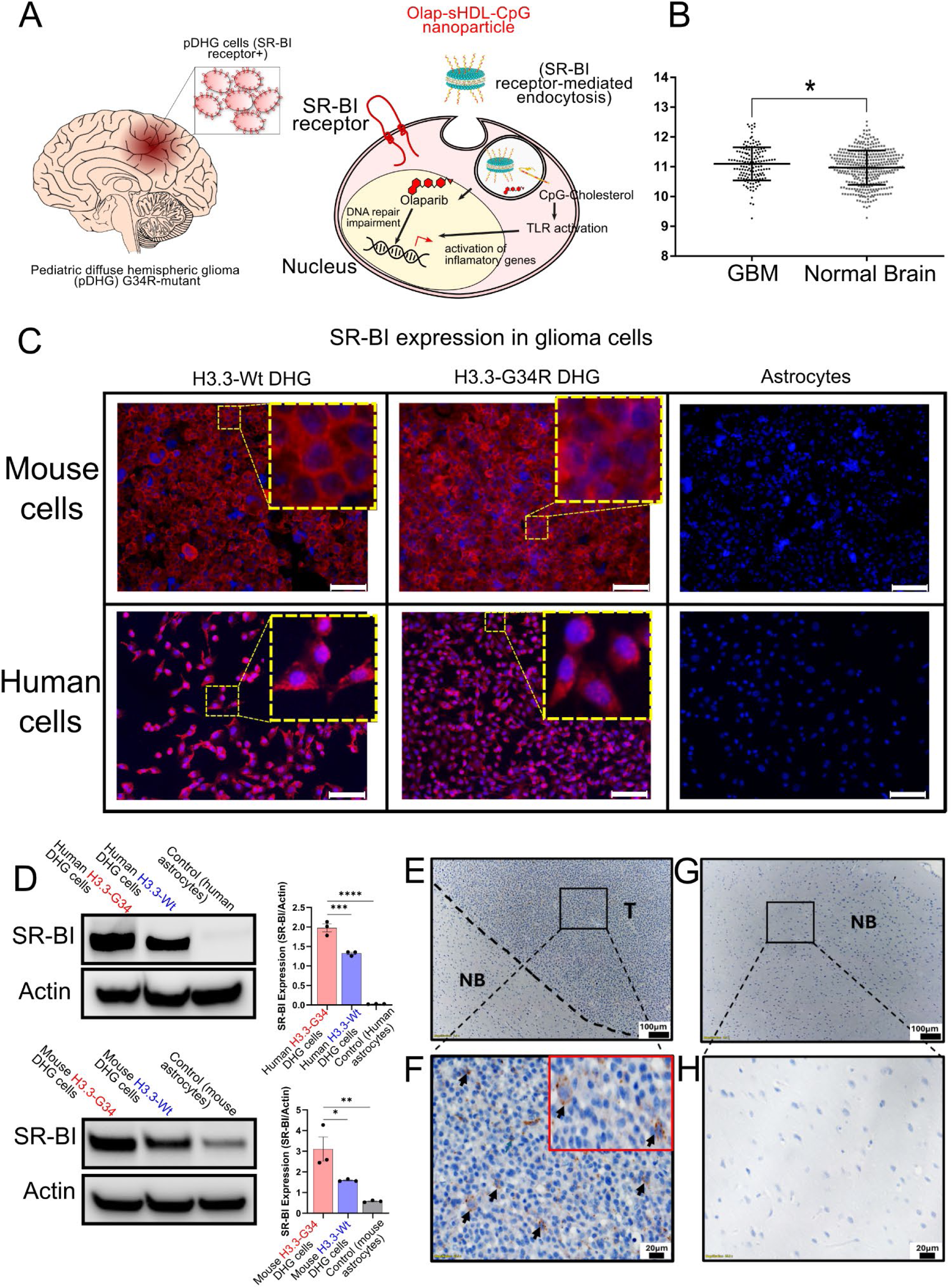
Expression of the HDL receptor SR-Bl in glioma cells. A) Illustration depicting the role of the SR-BI receptor in the uptake of sHDL nanoparticles. B) Violin plot showing increased SR-BI in GBM when compared with normal brain. *P<0.05. Mann-Withney test. C) Immunofluorescence of both mouse and human H3.3WT-DHG cells, H3.3-G34R DHG cells, and astrocytes expressing SR-B1 (red) and nuclei (blue). White scale bar represents 50µm. D) Western blot results showing SR-BI expression in both mouse and human H3.3-WT-DHG cells, H3.3-G34R-DHG cells, and astrocytes. E) Immunohistochemical detection of SR-BI expression in brain sections Representative IHC image at 10× magnification showing the tumor (T) and the normal brain (NB) separated by a dotted line (E). F) IHC image at 40× magnification showing strong SR-BI expression (brown staining) in tumor tissue with the inset showing the cells stained. (G) Normal brain structure at 10x magnification in the contralateral hemisphere. H) In contrast, no detectable SR-BI staining was observed in the adjacent normal brain region. Scale bar = 20 μm.

### Epigenetic and transcriptional activation of the NF-κB pathway in G34-mutant DHG cells

Given that we recently described an increase in the immune activation of G34-mutant DHG ^[16,20]^, we evaluated the expression of the NF-κB pathway genes in G34-mutant DHG cells. The NF-κB pathway is involved in the transcriptional activation of immuno-stimulatory cytokines which mediate the activation of the immune cells in the tumor microenvironment (TME) ^[27]^ (Figure 3A).

**Figure 3.**
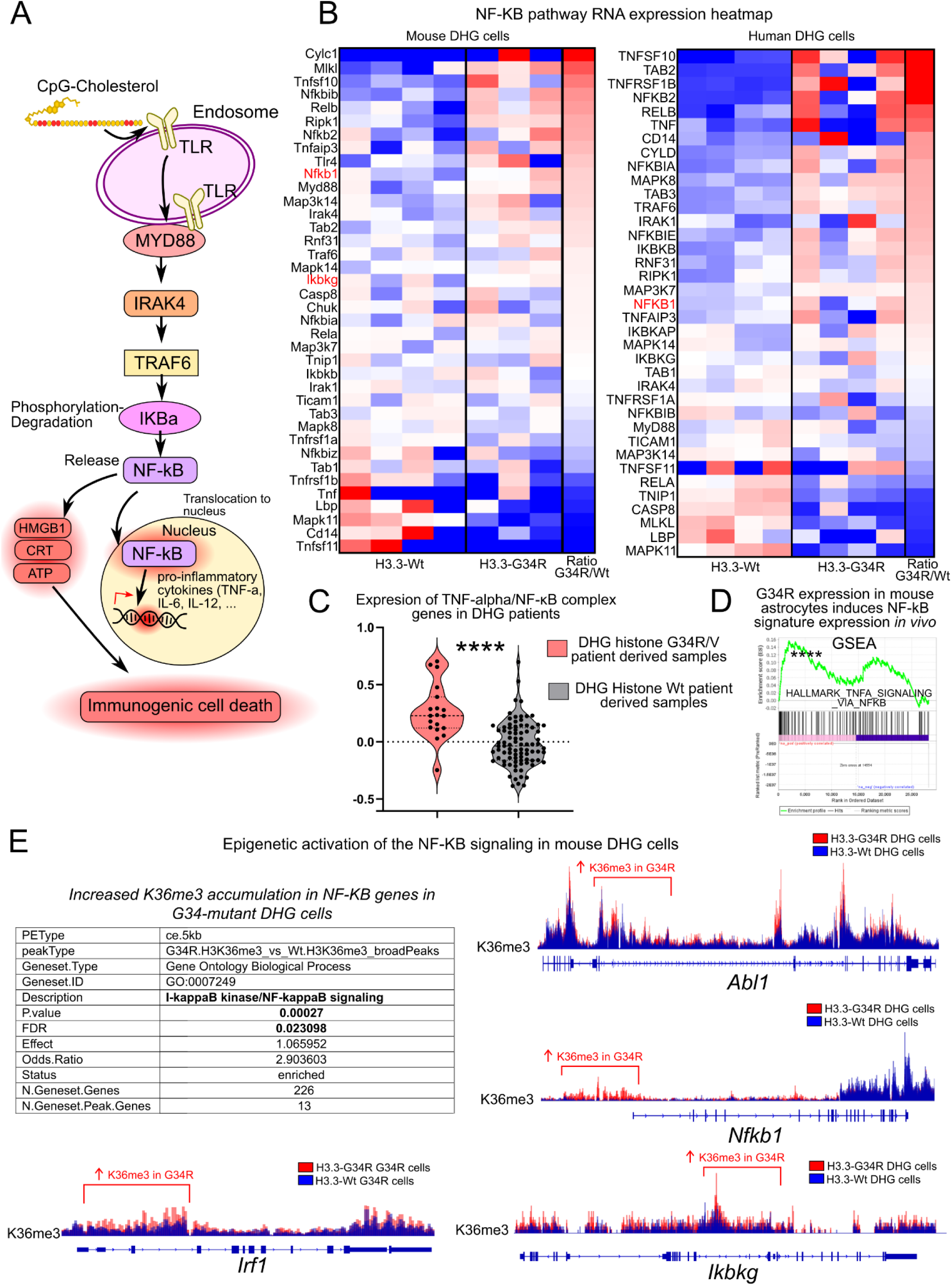
Epigenetic and transcriptional upregulation of the NF-KB pathway in G34-mutant DHG cells. A) Scheme depicting CpG-mediated activation of the NF-κB (Nuclear factor kappa-light-chain-enhancer of activated B cells) signaling. B) Transcriptional upregulation of NF-κB in mouse and human G34-mutant versus histone wild-type cells. C) Signature score of genes belonging to the TNF-alpha/ NF-κB complex in G34R/V-mutant and histone Wt DHG patient tumors samples. D) Expression of NF-κB signature in mouse astrocytes expressing histone-G34R versus Wt histone (FDR < 0.001). E) Epigenetic activation of NF-κB signaling genes in mouse G34-mutant versus histone wild-type cells and coverage plots of H3K36me3 in NF-κB complex genes in mouse G34-mutant versus histone wild-type cells.

Our results demonstrate that the NF-κB pathway is activated in human DHG genetically engineered to express H3.3-G34R compared to H3.3-Wt cells (Figure 3B) and in mouse H3.3-G34R DHG cells extracted from tumors (*in vivo*) compared to mouse H3.3-G34R DHG cells (Figure 3B) (Figure S1). We compared the expression of genes encoding proteins that comprise the NF-κB complex in histone-G34 mutant versus histone-Wt DHG patient tumors ^[6]^ observing significantly higher expression in the histone-G34 mutant DHG patients (Figure 3C). Moreover, we observed that the expression of G34R-mutant H3.3 histone in astrocytes in genetically engineered mouse derives in significantly higher NF-κB gene signature activation (Figure 3D).

To evaluate the correlation between the transcriptional upregulation of the NF-κB pathway genes with the activated epigenetic status of the genes belonging to this pathway in G34-mutant cells, we analyzed ChIP-seq data performed in mouse DHG cells and observed an increase in H3K36me3 levels in genes belonging to the pathway in G34-mutant cells. This indicates that the differences in gene expression in the NF-κB pathway genes are attributable to the presence of differential histone marks which are consequence of the expression of the G34-mutant histone (Figure 3E).

### Activation of the NF-κB pathway upon CpG stimulation in G34-mutant DHG cells

We evaluated the effect of the CpG treatment on the NF-κB pathway activity in the G34-mutant DHG cells *in vitro*. We incubated mouse DHG cells with sHDL-CpG nanoparticles or blank sHDL nanoparticles and after 24 hours of incubation, we collected total protein samples to analyze the activation of the NF-κB pathway with a posttranslational array features 215 antibodies related to the NF-kappa B pathway. The array has antibody spots that allow quantifying the total protein levels and the levels of the relevant post-translational modifications. Our results indicate that the total levels on post-translational marks that are associated with NF-κB pathway activation are upregulated upon treatment with sHDL-CpG nanoparticles versus blank sHDL nanoparticles, indicating that the CpG deoxynucleotides can activate this pathway in G34-mutant DHG cells (Figure 4A) (Figure S2). Specifically, posttranslational marks within the NF-κB complex were upregulated upon sHDL-CpG nanoparticles incubation (Figure 4B).

**Figure 4.**
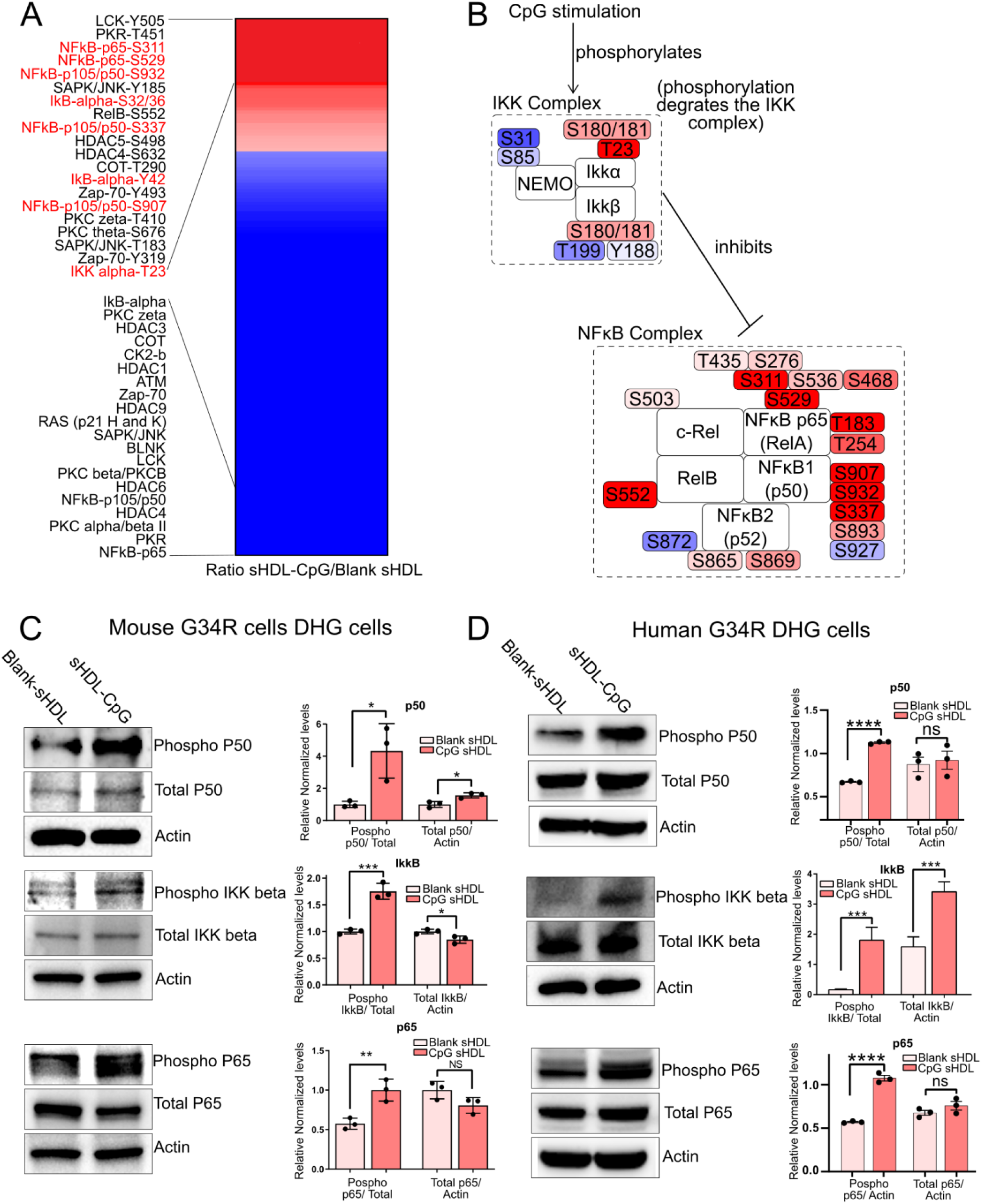
Activation of the NF-KB pathway in G34R DHG cells upon stimulation with sHDL-CpG nanoparticles. A) Protein phosphorylation array of NF-κB-related proteins in G34-mutant human DHG cells treated with sHDL-CpG nanoparticles versus sHDL blank nanoparticles. B) IKK and NF-κB complexes phosphorylation levels in G34-mutant human DHG cells treated with sHDL-CpG nanoparticles versus sHDL blank nanoparticles. C-D) Western blots of IKK and NF-κB complexes proteins in G34-mutant human (C) and mouse (D) DHG cells treated with sHDL-CpG nanoparticles versus sHDL blank nanoparticles.

To validate these results, we performed western blot to measure the phosphorylation of proteins belonging to the NF-κB complex (P50 and P65) and to the IKK complex (the main inhibitory complex that modulates the NF-κB complex activity (Figure 4C-D). The inhibitory activity of the IKK complex is blocked by phosphorylation marks that result from transduction signals, among them, TLR9 activation). Both mouse and human G34-mutant cells have higher levels of phospho-P50, phospho-P65 and phospho-IKK-beta upon stimulation with sHDL-CpG nanoparticles in comparison with blank sHDL nanoparticles (Figure 4C-D).

These results indicate that the CpG nanoparticles can be used to modulate the NF-κB activity in G34R-mutant DHG.

### Analysis of the cell cycle and DNA repair signaling on G34-mutant DHG cells upon stimulation with olaparib-sHDL-CpG nanoparticles

To evaluate the effect of the PARP inhibitor olaparib incorporated the sHDL nanoparticles, we incubated mouse DHG cells with, olaparib-sHDL-CpG nanoparticles or blank sHDL nanoparticles and after 24 hours of incubation, we collected total protein samples to analyze the activation of the cell cycle and DNA repair signaling with a posttranslational array features 238 antibodies important to the cell cycle control system and DNA damage/repair mechanism. Our results show increase in PTMs that indicate the activation of DNA repair processes BRCA1 (Phospho-Ser1524), RAD52 (Phospho-Tyr104), BRCA1 (Phospho-Ser1423), DNA-PK (Phospho-Thr2638) (Figure S3).

We also observed an increase in PTMs that indicate the induction of p53-mediated apoptosis, p53 (Phospho-Ser378, Phospho-Ser20, and Phospho-Ser9) (Phosphorylation at Ser20 stabilizes p53 by disrupting its interaction with MDM2, an E3 ubiquitin ligase that targets p53 for degradation; phosphorylation at Ser378 modulates p53’s transcriptional activity, promoting apoptosis; and decreased phosphorylation at Ser9 may also influence p53’s ability to induce apoptosis. Therefore, these changes in p53 phosphorylation suggest that PARPi treatment might be promoting apoptosis (Figure S3).

Finally, our results show increased levels of CDC25C (Phospho-Ser216) and Cyclin D1 (Phospho-Thr286). CDC25C phosphorylation at Ser216 creates a 14-3-3 binding site that sequesters CDC25C in the cytoplasm, preventing it from dephosphorylating and activating the Cyclin B–CDK1 complex required for G₂/M transition. Similarly, phosphorylation of Cyclin D1 at Thr286 promotes its nuclear export and subsequent ubiquitin-mediated degradation, thereby reducing Cyclin D1–CDK4/6 activity and enforcing a G₁ arrest. Together, these modifications indicate that olaparib-sHDL-CpG treatment not only triggers DNA repair and p53-mediated apoptosis but also induces activation of cell cycle checkpoints to halt tumor cell proliferation.

### NF-κB Inhibitors reduce the efficacy of olaparib-sHDL-CpG nanoparticles

To corroborate the role of NF-κB in the therapeutic effect of the olaparib-sHDL-CpG nanoparticles, we studied the cytotoxicity of the nanoparticles in the presence of NF-κB inhibitors. Our results show that both two NF-κB Inhibitors, JSH-23, a drug that specifically targets the nuclear translocation of the p65 subunit and its transcriptional activity, and PDTC, a compound that blocks the phosphorylation of IκB (Figure 5A), reduce the efficacy of olaparib-sHDL-CpG nanoparticles on mouse and human G34-mutant DHG cells (Figure 5B-C).

**Figure 5.**
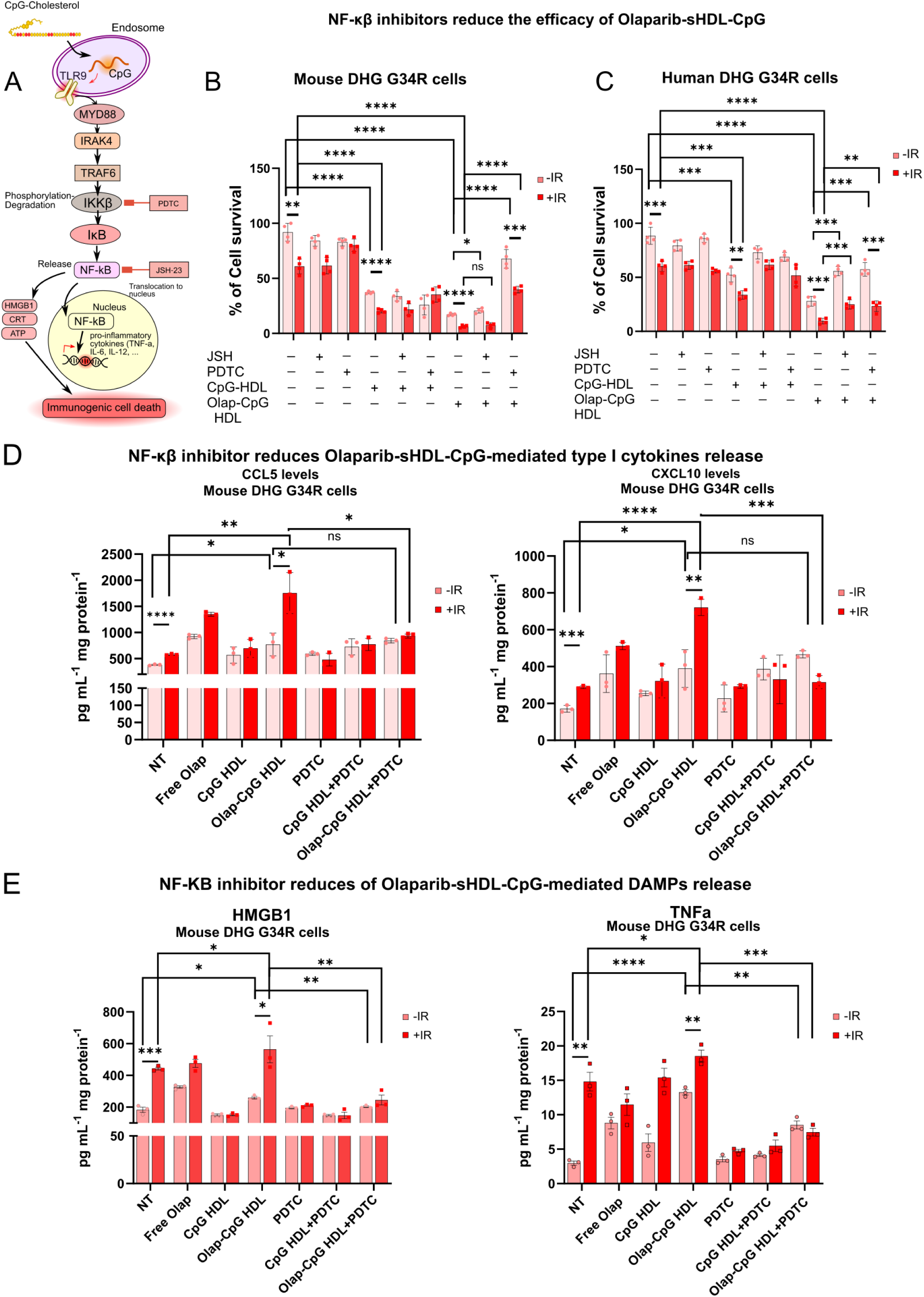
NF-κβ inhibitors abrogate Olap-CpG-HDL-induced cytotoxicity and radiosensitivity in mouse and human H3.3-G34R glioma cells. A) Scheme depicting the effect of NF-κβ inhibitors PDTC and JSH-23 on NF-κβ pathway activation. B) Effect of NF-κβ inhibitors on sHDL-CpG and Olaparib-sHDL-CpG nanoparticles-mediated cytotoxicity in mouse DHG G34R cells. C) Effect of NF-κβ inhibitors on sHDL-CpG and Olaparib-sHDL-CpG nanoparticles cytotoxicity in human DHG G34R cells. D) Effect of NF-κβ inhibitors on sHDL-CpG and Olaparib-sHDL-CpG nanoparticles-mediated cytokine release in mouse and human DHG G34R cells. E) Effect of NF-κβ inhibitors on sHDL-CpG and Olaparib-sHDL-CpG nanoparticles-mediated DAMPs release in mouse and human DHG G34R cells.

Inhibition of NF-κB signaling with PDTC also blocks the release of cytokines and DAMPS in G34-mutant DHG cells (Figure 5D-E).

### Olaparib-sHDL-CpG nanoparticles show increased efficacy *in vivo* compared to free olaparib

To assess the response of H3.3-G34R DHG to olaparib-sHDL-CpG nanoparticles *in vivo*, H3.3-G34R DHG mouse cells were implanted into the striatum of mouse Wt mice (C57BL/6).

G34-mutant DHG–bearing mice were treated H3.3-G34R mice at 14 days, after implantation, with free olaparib, empty sHDL nanoparticles, olaparib-sHDL-CpG nanoparticles in combination with radiation therapy (RT), for two weeks (Figure 6A). Empty sHDL nanoparticles without radiation were used as treatment negative control (vehicle control).

**Figure 6:**
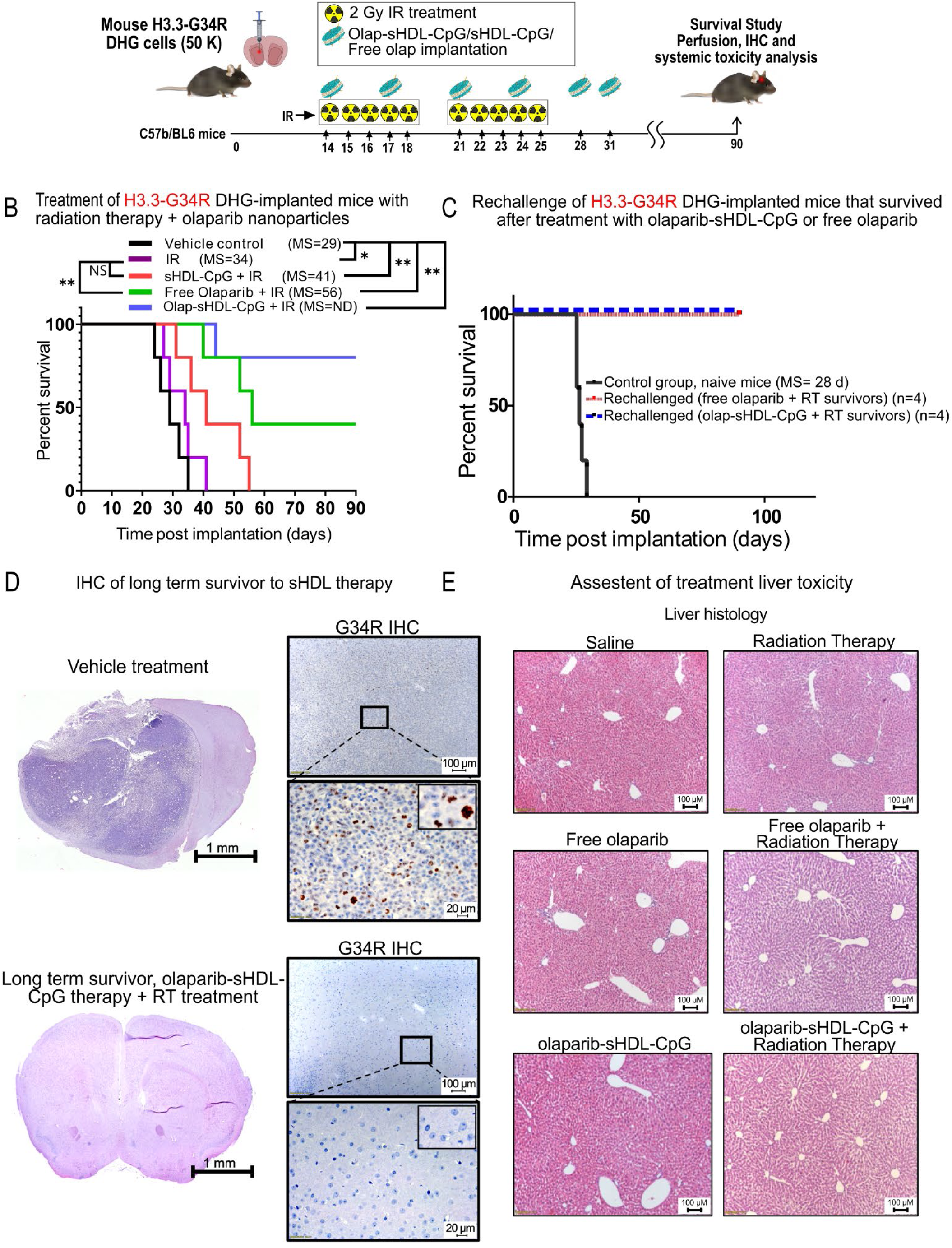
Survival of mice bearing G34-mutant DHG upon Olaparib-sHDL-CpG therapy. Efficacy of Olaparib-sHDL-CpG in intracranial H3.3-Wt DHG. (A) Scheme depicting the preclinical design to test the impact of Olaparib-sHDL-CpG vs. free Olaparib combined with RT in mice bearing H3.3-G34R pediatric diffuse hemispheric glioma. B) Kaplan-Meier survival curve of mice bearing H3.3-G34R pediatric diffuse hemispheric gliomas treated according to treatment groups indicated. Curves were compared using log-rank test with *P<0.05; **P<0.01 (n = 5 mice per experimental group). C) Kaplan-Meier survival plot of H3.3-G34R–bearing mice that survived following Olaparib-sHDL-CpG + RT or free olaparib + RT therapies and that were rechallenged with H3.3-G34R DHG cells, compared with naive mice implanted with the same cells (control group). D) H&E and G34R-mutation IHC image of vehicle (empty sHDL) treated and a long-term survivor to Olaparib-sHDL-CpG therapy + RT treatment mouse brains. E) Histology analysis of livers of mice from different treatment groups. *P < 0.05, **P < 0.01; log-rank (Mantel-Cox) test.

All the animals in vehicle control and only radiation therapy (RT) groups reached an endpoint, with median survivals of 29 and 41 days respectively. Forty percent of the animals in the free olaparib + RT group survived versus 80% in the olaparib-sHDL-CpG nanoparticles + RT group, with a median survival of 56 days for free the free olaparib + RT group and not reached MS for the olaparib-sHDL-CpG nanoparticles + RT group. Kaplan-Meier analysis indicates statistical significance for olaparib-sHDL-CpG nanoparticles + RT versus free olaparib + RT (p <0.05) (Figure 6B). The treatment performed in absence of radiation therapy did not show differences between the olaparib-sHDL-CpG nanoparticles and the empty particles or free drug, indicating that DNA damage is necessary to reach therapeutic efficiency of the Olaparib + CpG combination (Supplemental Figure 6).

Surviving mice of the free olaparib + RT and olaparib-sHDL-CpG nanoparticles + RT groups were rechallenged with a second implantation of G34-mutant mouse DHG cells in the contralateral hemisphere, and none of the animals developed tumors, whereas 100% of the naïve control animals developed tumors (Figure 6C). This indicates that the mice that survived following the free olaparib + RT and olaparib-sHDL-CpG nanoparticles + RT treatments developed antitumor immunological memory.

To assess the safety of the treatment, we analyzed its toxicity by evaluating brain architecture (Figure 6D), liver histology (Figure 6E), complete blood counts (CBCs), and on surviving mice treated with olaparib-sHDL-CpG nanoparticles + RT (Supplemental Figure 7). Liver histology revealed no abnormalities in livers comparing nontreated (NT) animals and mice treated with olaparib-sHDL-CpG nanoparticles + RT. No significant changes in CBC levels were found untreated and treated mice. Altogether, this data indicates no evidence of toxicity due to the treatment.

Treatment was finally assessed by IHC on treated tumor-bearing brains of each experimental treatment (Supplemental Figures 8 and 9). Long term survivors of the olaparib-sHDL-CpG nanoparticles + RT group do not show signs of brain damage (MBP unspecific staining), no signs of inflammation (low GFAP) and absence of activated microglia and macrophages (CD68-) and absence of residual tumor cells (G34R-); whereas the control-treated animals and the animals treated with RT and free olaparib that showed tumor showed more intense MBP, GFAP, CD68 and G34R staining.

## Materials and Methods

### Preparation and characterization of sHDL nanoparticles

20 mg DMPC, 10 mg 22A, and 1 mg Olaparib were dissolved in acetic acid and mixed. Mix was frozen-dry in liquid nitrogen for 24 h. The resulting powder was resuspended in ∼0.9 mL of pH 7.4, 10 mM phosphate buffer was added to the freeze-dried powder, and the mix was heated and cooled in a 50 °C water bath and ice bath for 5 min each for three cycles. Mix was bath-sonicated for 3 min at the lowest power. The pH of the resulting Olaparib-sHDL-CpG nanodisc solution was adjusted to 7.4, and 1.5 mg CpG-Chol were incorporated into the nanodisc (final CpG concentration of 75 ug/mL) by incubating for 2 h at room temperature. The olaparib concentration in the final formulation was adjusted to is 1 mg/mL.

The mobile phases A, B, and C were delivered at 0.3 mL min⁻¹ and consisted of 0.1% formic acid in water, acetonitrile, and methanol, respectively. Incorporation efficiency was determined as the ratio of the olaparib concentration after purification to that measured before purification. Incorporation of chol-CpG into the nanodiscs was confirmed by size-exclusion chromatography on a Waters 1525 HPLC system (Waters, Milford, MA) equipped with a Tosoh TSKgel G3000SWXL 7.8 mm × 30 cm column (Tosoh Bioscience, King of Prussia, PA), using PBS as the mobile phase at a flow rate of 1 mL min⁻¹.

### Particle size

The hydrodynamic diameter of the sHDL nanodiscs was measured by dynamic light scattering (DLS) on a Malvern Zetasizer after diluting each formulation in PBS to 1 mg mL⁻¹ (22A peptide).

#### Electron microscopy

Particle morphology was assessed by transmission electron microscopy (TEM) with negative staining. sHDL samples were diluted in buffer to 10 μg mL⁻¹ (22A peptide), and 4 μL was applied for 1 min to a glow-discharged 400-mesh copper grid with formvar/carbon film (Electron Microscopy Sciences). Grids were washed and negatively stained with 0.1% uranyl acetate, then imaged on an FEI Morgagni electron microscope operated at 100 kV and 22 000× magnification.

### Genetically engineered mouse glioma models

The development of the H3.3-G34R and H3.3-Wt DHG mouse models was described in ^[16]^. The cells were maintained in neural stem cell medium (DMEM/F12 with l-glutamine, Gibco, Thermo Fisher Scientific, catalog 11320-033), B-27 supplement (1×) (Gibco, Thermo Fisher Scientific, catalog 12587-010), N-2 supplement (1×) (Gibco, Thermo Fisher Scientific, catalog 17502-048), penicillin-streptomycin (100 U/mL) (Cellgro, catalog 30-001-CI), and normocin (1X) (InvivoGen, catalog ant-nr-1) at 37°C, 5% CO2. FGF and EGF (Shenandoah Biotech, catalogs 100-26 and 100-146) were added twice weekly at a concentration of 1 μL (20 ng/μL each stock, 1,000× formulation) per 1 mL medium. Cells were verified for GFP and Katushka expression by flow cytometry. Five cell cultures generated independently for each phenotype (H3.3-WT and H3.3-G34R mouse DHG) were used for the experiments.

### Genetically engineered human glioma models

The development of the H3.3-G34R and H3.3-Wt DHG SJ-GBM2-derived human models was described in ^[16]^. Briefly, GEMM cells were generated by stable transfection of SJ-GBM2 with plasmids expressing H3.3-G34R or H3.3-Wt. SJ-GBM2 cells (CVCL_M141) were a gift of the COG Repository at the Health Science Center, Texas Tech University (Lubbock, Texas, USA). SJ-GBM2 cells were grown in IMDM (Gibco, Thermo Fisher Scientific, catalog 1244005320) supplemented with 20% FBS at 37.0°C, 5% CO2, 20% O2 according to a previously published report ^[27]^. Cells were used in early passages and tested regularly for mycoplasma and G34R or H3.3 expression. Three independent polyclonal populations were obtained for each genotype (SJ-GBM-2-H3.3-G34R and SJ-GBM-2-H3.3-WT). These cells are referred to as H3.3-G34R and human DHG cells.

### Dose-response curve assessment of olaparib-sHDL-CpG nanoparticles

We assessed the susceptibility of H3.3-G34R and H3.3-Wt mouse and human cells to unformulated olaparib and olaparib-sHDL-CpG nanoparticles. 2×10^3^ mouse or human cells per well on 96-well plates in 190 µL of media were seeded in 190 µL of media. We used quintuplicated wells per inhibitor dose evaluated for each cell type. We evaluated seven concentrations of inhibitors in serial logarithmic dilutions (e.g., 1 µM, 3 µM, 10 µM, 30 µM, 100 µM and 300 µM). One day after plating the cells 10 µL of 20X concentrated inhibitor was added to each well for each dilution evaluated. After three days of incubation, viability was assessed with CellTiter Glo assay (Promega, Cat. # G7572). The results were analyzed in GraphPad Prism 7 using a sigmoidal regression model, which allows the estimation of the IC50 and evaluation of the statistical differences among dose-response curves.

### Immunofluorescence analysis

For human SJGBM-derived DHG cells, which are adherent, we placed sterilized round shaped coverslips into 24-well plates and seeded 2×10^4^ cells per well in 0.5 mL of media. For floating cells (mouse primary DHG cells), 2×10^5^ cells per well in 6-well plates in two mL of media were plated. Cells were incubated for 24 hours, and, for adherent cells, cells were washed twice with PBS and fixed with 300-400 µL of buffered 3% formaldehyde for each well and cells were incubated at room temperature for 20 minutes. For floating cells, the cells were separated by incubation with Accutase and after washing, the cells were resuspended and loaded into a Cytospin cassette and were centrifuged at 500 RPM for ten minutes in Cytocentrifuge (Thermo, Cytospin 4) to deposit the cells in a slide. Slides were washed twice with PBS incubated with buffered 3% Formaldehyde fixative solution on top of each slide at room temperature for 20 minutes. The subsequent steps of the protocol are the same for adherent and floating cells. Wells/slides were washed twice with PBS and twice with 400 µL of wash buffer (0.1% BSA in 1X PBS). Next, cells wells/slides were blocked for non-specific staining with 400 µL of blocking buffer (10% normal horse serum, 0.3% Triton X-100) for 45 minutes at room temperature. Following blocking, samples were incubated with primary antibody prepared in dilution buffer (1% BSA, 1% normal horse serum, 0.3% Triton) for one hour at room temperature. Wells/slides were washed twice with 400 µL of wash buffer (0.1% BSA in 1X PBS) and incubated for one hour at room temperature in dark with fluorescent-conjugated secondary antibodies. After, wells/slides were washed twice with 400 µL of wash buffer (0.1% BSA in 1X PBS) and stained with 300 µL DAPI solution (1:5000 of a 14.3 mM DAPI stock solution) for two minutes, rinsed with PBS and mounted with ProLong™ Gold Antifade Mountant (ThermoFisher, Cat. # P36930). Slides were preserved at 4°C and imaged after 24 hours using an Olympus BX53 microscope and cellSens Dimension software (Olympus).

### Western blotting

For each experimental point, mouse and human H3.3-Wt and H3.3-G34R DHG cells were seeded at a density of 3 x 10^6^ cells into 75-cm^2^ flasks. Cells were incubated for 24 hours to allow attachment; and subjected to treatment. Cells were washed three times with PBS and cell lysates were generated by resuspension in 0.2 mL RIPA lysis buffer (ThermoFisher, Catalog # 89900) containing protease inhibitors (ThermoFisher, Halt™ Protease and Phosphatase Inhibitor Cocktail (100X), Catalog # PI78440) on ice for five minutes. Cell lysates were centrifuged at (13,000 RPM, 10’, 4°C) to remove unsuspended debris and supernatants were collected. Protein concertation was determined via bicinchoninic acid assay (BCA) (Assay Kit Pierce BCA, Cat # PI23227, Thermo Scientific). For electrophoretic separation of proteins, 20 µg of total protein were resuspended in loading buffer (10% sodium dodecyl sulfate Sigma Aldrich 71736, 20% glycerol, and 0.1% bromophenol blue) incubated five minutes at 95°C and loaded onto a 4-12% Bis-Tris gel. Proteins from the gel were transferred to 0.2 µm nitrocellulose membrane and blocked with 5% nonfat milk in TBS-0.1% Tween-20. After blocking, membranes were incubated with primary antibodies overnight at 4°C with gentle movement. The next day, blots were washed with TBS-0.1% Tween-20 and incubated with secondary (1:4000) antibody for one hour at room temperature. Blots were washed several times again with TBS-0.1% tween-20 and visualized under Biorad gel imaging software. Band intensities were quantified using GelAnalyzer software. A list of the antibodies used for WB and other experiments is provided in Table S1.

### Post-translational modifications Phospho-Arrays

Post-translational modifications were evaluated using phospho-antibody arrays. To measure phosphorylation-dependent activation of proteins involved in DNA repair and cell-cycle control, we used two slide-based antibody arrays from Full Moon BioSystems: the NFκB Phospho Antibody Array (Cat. # PNK215; 215 site- and phospho-specific antibodies) and the Cell Cycle Control Phospho Antibody Array (Cat. # PCC238; 238 site-and phospho-specific antibodies) covering components of the NFκB pathway, cell-cycle regulation, and DNA damage response/repair.

For the NFκB array, mouse DHG cells (2 × 10^6^) were incubated with sHDL-CpG nanoparticles or blank sHDL nanoparticles. After 24 h, total protein lysates were collected and processed according to the Antibody Array Assay Kit protocol (Full Moon BioSystems, Cat. # KAS02). For the Cell Cycle Control array, mouse DHG cells were treated with olaparib-sHDL-CpG nanoparticles or blank sHDL nanoparticles for 24 h, then harvested for total protein to assess cell-cycle and DNA repair signaling.

Following lysate preparation, protein concentrations were determined by BCA assay (Pierce BCA Assay Kit, Thermo Scientific, Cat. # PI23227) according to the manufacturer’s instructions. For each condition, 100 µg of protein was biotinylated using the KAS02 kit protocol. Array slides were blocked, then incubated with the biotinylated samples to allow binding to the immobilized antibodies. After washing, bound proteins were detected with Cy3-streptavidin, and slides were scanned using a GenePix 4100A Microarray Scanner (UMICH Single Cell Analysis Resource Core).

Images were processed in ImageJ ^[28]^, and spot intensities were extracted using the Protein Array Analyzer plugin ^[29]^. Signal was first normalized to the average actin intensity on each array. For each phospho-specific target, the normalized phospho signal was additionally normalized to the corresponding total-protein antibody signal. Each target is represented by six replicate spots; therefore, six normalized intensity values were obtained per site-specific/phospho-specific measurement. Statistical analyses were performed in GraphPad Prism 7 using unpaired t-tests to compare treated versus untreated conditions for each protein and phospho-site. Fold changes and multiple-testing–adjusted significance values (false discovery rates) were reported for each marker.

### Cell survival assay

Sleeping-beauty derived genetically engineered mouse models i.e. H3.3wt (NRAS/shP53/shATRX) and H3.3-G34R (NRAS/shP53/shATRX/H3.3G34R) neurospheres, and human genetically engineered pediatric diffuse hemispheric glioma (DHG) cells i.e. SJ-GBM2 H3.3-Wt and H3.3G34R cells, were plated at a density of 1000 cells per well in a 96-well plate (Fisher, 12-566-00) 24 hours prior to treatment. Cells were then treated with one of the following: saline, free-CpG (ODN 1826; InvivoGen, tlrl-1826), empty-HDLs, sHDL-CpGs, free-olaparib (at IC50 doses, AZD2281; Selleckchem, S1060), free-JSH23 (10µM; NF-κβ transcriptional inhibitor [MedChemExpress; HY-13982], inhibits nuclear translocation of NF-κβ p65 without affecting Iκβα degradation), free-PDTC (5µM; Ammonium pyrrolidine dithiocarbamate (PDTC), NF-κβ functional inhibitor [Selleckchem; S3633] inhibits NF-κβ by inhibiting IκB phosphorylation, thus blocking NF-κβ translocation to the nucleus, reducing the expression of downstream cytokines), and olaparib-CpG loaded-HDLs (i.e. olaparib-sHDL-CpGs, at equivalent IC50 doses of olaparib), either alone or in combination with radiation for 72 hours in triplicate wells per condition. Cells were pretreated with 2h prior to irradiation with 3 Gy and 6 Gy of radiation, for mouse neurospheres and human DHG cells respectively. sHDLs without olaparib-CpGs were tested in the same peptide and lipid concentrations as olaparib-sHDL-CpG with specified olaparib concentration (IC50). Cell viability was determined with CellTiter-Glo 2.0 Luminescent Cell Viability Assay (Promega, G9242) following manufacturer’s protocol. The resulting luminescence was read with the Enspire Multimodal Plate Reader (PerkinElmer). Data were represented graphically using the GraphPad Prism 9.0 software, and statistical significance was determined by one-way ANOVA followed by Tukey’s test for multiple comparisons.

### Measurements of DAMPs and Type I Interferons in the tumor conditioned media

To analyze the levels of DAMPs, type I interferons, and inflammatory cytokines in the tumor-conditioned media, genetically engineered H3.3-Wt and H3.3-G34R mouse neurospheres were seeded at a density of 1 x 10^6^ cells into a 6-well plate. The cells were then allowed to settle overnight before treatment. On the next day, cells were treated with either saline, empty-sHDLs, sHDL-CpGs, free-olaparib (at IC50 doses), free-PDTC (at 5µM doses; Ammonium pyrrolidine dithiocarbamate (PDTC) [Selleckchem; S3633] inhibits NF-κβ by inhibiting Iκβ phosphorylation, thus blocking NF-κβ translocation to the nucleus, reducing the expression of downstream cytokines), and olaparib-CpG loaded-HDLs (olaparib-sHDL-CpGs, at equivalent IC50 doses of olaparib) for 2 hours in triplicate wells prior to receiving 3Gy (for H3.3-Wt and H3.3-G34R mouse neurospheres) of radiation. sHDLs without olaparib-CpGs were tested in the same peptide and lipid concentrations as olaparib-sHDL-CpG with specified olaparib concentration (IC50). Release of DAMPs was assessed 72 hours post-irradiation. Concentrations of HMGB1, IL33, IL6, IL1α, TNF-α, CCL2, CCL5, CCL4, CXCL1, CXCL2, CXCL5, CXCL10, IL18, IL2,

IL12p70 in the culture supernatants were measured by quantitative ELISA, either following the manufacturer’s protocol (Mouse HMGB1/HMG-1 ELISA Kit; Novus Biologicals, NBP2-62767) or through Immunology Core at the Rogel Cancer Center, Michigan Medical School. All the cytokine levels were normalized by respective protein concentrations.

### Primary mouse Astrocytic culture

Cortices from P0-P4 C57/BL6 mice pups were isolated in ice-cold dissection media, consisting of 0.11 mg/mL sodium pyruvate, 0.1% glucose, 10 mM HEPES in 1x HBSS (Ca2+ and Mg2+ -free). This was followed by 0.25% trypsin digestion for 20 min at 37°C. The tissue was then triturated 10-15 times with pipette tips of varying size. The filtered cell suspension was then centrifuged for 6 minutes at 300g x to pellet cortex tissue pieces. The supernatant was removed, and the cells were resuspended in Glial media consisting of Dulbecco’s Modified Eagle Medium (DMEM) with 10% Fetal Bovine Serum (FBS), 0.1% glucose and 100 U/mL penicillin/streptomycin. The cell suspension was added to the Glial media at a ratio of 1:10 and added to a T75 flask, which was coated with 20 ml of poly-D-lysine (Thermo Fisher Scientific, A3890401) at a concentration of 50 μg/ml in cell culture grade water for 1 hour at 37 °C in the CO2 incubator, a day prior to the dissection.

On day 3 in vitro, the flask was shaken vigorously to separate astrocytes from the other types of brain cells (OPCs, neurons, ependymal and microglial cells). This can be done because, unlike other brain cells, astrocytes maintain adherence to the bottom of the flask. Once the astrocytes were around 70%-80% confluent, they were split and shaken again until they were pure astrocytes without any other types of cell contamination. Total protein extracts were prepared from normal mouse astrocyte cells in a RIPA lysis extraction buffer (Thermo Fisher Scientific, Pierce, 89900) with 1X of Halt protease and phosphatase inhibitor cocktail (Thermo Fisher Scientific, 78442).

### Tumor implantation

Mice were anesthetized with intraperitoneal injections of ketamine (75 mg/kg) and dexmedetomidine (0.5 mg/kg) prior to stereotactic implantation. A total of 50000 H3.3G34R DHG neurospheres were injected intracranially at 0.5 mm anterior and 2.0 mm lateral from the bregma, and 3.0 mm ventral from the dura. Neurospheres were injected at a rate of 1 μL/min. Post-surgical analgesia included buprenorphine (0.1 mg/kg, subcutaneously) and carprofen (5 mg/kg, subcutaneously). 14 days post-implantation, mice were randomly divided into groups for different treatments, including: vehicle control, sHDL-olap, irradiation, free olaparib, empty sHDL+IR, free olaparib +IR, and olaparib-sHDL-CpG + IR (2 Gy/dose, total 20 Gy). The long-term survivors from the treatment were rechallenged in the contralateral hemisphere using the same coordinates, 0.5 mm anterior and -2.0 mm lateral from the bregma, and -3.0 mm ventral from the dura. At the moribund stage, mice were perfused with Tyrode’s solution and PFA. The brains and livers were collected from these mice and embedded for immunohistochemistry. All procedures involving mice were performed following policies set by the Association for Assessment and Accreditation of Laboratory Animal Care (AAALAC) and the Institutional Animal Care and Use Committee (IACUC) at the University of Michigan (PRO00011290).

### *In vivo* irradiation

Mice assigned to radiation treatment groups underwent fractionated ionizing radiation starting two weeks after tumor implantation. Mice were subjected to an irradiation (IR) dose of 2 Gy for 5 days a week for 2 weeks for a total of 20 Gy of ionizing radiation. The procedure was performed as follows: Briefly, mice were lightly anaesthetized with isoflurane. Mice were then placed under a copper orthovoltage source, with the irradiation beam directed to the brain and body shielded by iron collimators. Irradiation treatment was given to mice at the University of Michigan Radiation Oncology Core.

### Immunohistochemistry

Tissues from the brains were first fixed in 4% paraformaldehyde (PFA) and embedded in paraffin. 5 μm thick sections were cut using (Leica RM2165) microtome system. Sections were permeabilized in TBS containing 0.5% Triton-X (TBS-Tx) for 20 min, which was followed by heat induced antigen retrieval at 96 °C with 10 mM sodium citrate (pH 6) for 20 min. The sections were cooled to room temperature (R.T.) and rinsed 5 times with TBS-Tx (5 min per wash). Samples were blocked with 5% goat serum for 2h at R.T. The sections were then incubated with primary antibody against GFAP (1:200), MBP (1:200), CD68 (1:1000), Anti-Iba1 (1:2000), Anti-H3.3-G34R (1:500), Anti-CD8 (1:100) and Anti-GR1 (1:50) and Anti-SRB1 (1:200) diluted in 1% goat serum and incubated overnight at 4 °C. The next day, sections were washed with TBS-Tx and incubated with biotin-conjugated secondary antibody (1:1000 in 1% goat serum TBS-Tx) for 4 hrs at R.T. These sections were subjected to 3, 3′- diaminobenzidine (DAB) (Biocare Medical) with nickel sulfate precipitation. The reaction was quenched with 10% sodium azide and washed in 0.1 M sodium acetate. Sections were dehydrated through xylene and mounted with DePeX medium (Electron Microscopy Sciences). Images at 10X and 40X magnification were obtained using brightfield microscopy (Olympus BX53). For histopathological assessment, paraffin-embedded 5 μm brain sections from each treatment group were stained with hematoxylin and eosin (H&E).

### Liver histochemistry

Livers were embedded in paraffin for histological analysis, 5 μm-thick sections were cut using the microtome system and sections were stained using H&E. Bright-field images were acquired employing an Olympus MA BX53 microscope.

### Serum Chemistry Analysis

Blood was taken from the submandibular vein from DHG glioma-bearing mice and transferred to serum separation tubes (Biotang). Samples were incubated at room temperature for 90 min to allow for blood coagulation. Tubes were then centrifuged at 3000 rpm (500 *g*). Complete serum chemistry for all samples was assessed by the Veterinary Core Facility at the Medical School.

## Discussion

Despite the breakthroughs in cancer treatment over the last decades, the benefit for pediatric diffuse hemispheric gliomas has been limited. The recent molecular classification of these tumors has been of vital importance to determine directions for treatments. For example, it indicated that pediatric gliomas differ significantly from their adult counterparts, and that therapeutic approaches designed for adult gliomas do not adapt well to pediatric gliomas. We have shown that DHG tumors harboring G34R/V mutations in the histone H3.3 exhibit decreased DNA repair functionality. This translates into specific vulnerability to therapies combining DNA damage (as radiation therapy) and DNA repair inhibitors, such as the PARP inhibitor olaparib. We further demonstrated that DNA damage combined with DDR inhibitors leads to activation of the cyclic GMP–AMP synthase (cGAS)–stimulator of interferon genes (STING) pathway, enhancing the immunological antitumoral response.

Despite these encouraging preliminary results, PARP inhibitor clinical trials have failed to demonstrate positive efficacy to date ^[30]^. The discordance between the molecular and preclinical results and clinical efficacy can be attributed to several factors, including different mechanisms of action among PARP inhibitors, lack or deficient molecular tumor stratification in the clinical studies, discrepancies in drug activity or median life between preclinical models and patients, and poor drug penetrance. We demonstrated before that Pamiparib, still untested in clinical trials, has potent bioactivity *in vivo* against G34R-mutant DHG mouse models ^[16]^. olaparib, although potent *in vitro*, has poor penetrance through the blood-brain barrier, making it a suboptimal choice for glioma treatment.

For this reason, we evaluate the therapeutical efficacy of olaparib formulated into sHDL nanoparticles. We showed before that formulation into sHDL nanoparticles enhances drug delivery to glioma tumors *in vivo* ^[23]^. We demonstrate in the present study that glioma cells exhibit upregulation of the sHDL receptor SR-BI in comparison with non-cancerous brain cells. This finding provides a molecular mechanism that supports our previous in vivo results ^[23]^, which showed a high level of specificity in sHDL nanoparticle uptake by glioma cells. Thus, sHDL nanoparticles represent an optimal delivery strategy to minimize off-target drug effects and maximize drug exposure in target cells.

To further exploit the DNA-damage mediated immune response specific to G34-mutant DHG, we combined the PARP inhibitor with CpG dinucleotides, which activate immune cells by triggering NF-κB activation.

We demonstrate that G34R mutant cells show higher basal levels of NF-κB activation. This might be associated with the genetic instability triggered by the expression of the G34 mutant histone, which promotes immune-activity in G34-mutant DHG ^[16,20]^. We demonstrate that NF-κB activation boosts the therapeutic efficacy of the PARP inhibitor olaparib against G34-mutant DHG *in vitro*, and that the effects of CpG in these cells are driven by NF-κB activation. We also showed that CpG elicits the release of cytokines and DAMPs in G34-mutant DHG cells, and that this effect is dependent upon NF-κB activation.

In summary, our approach exploits the DNA repair deficiency and immune-permissive microenvironment of G34-mutant DHG, using a dual-drug nanoparticle formulation that specifically targets glioma cells in combination with irradiation-induced DNA damage.

The evaluation of this therapeutic approach *in vivo* demonstrated improved survival compared to the administration of free olaparib *in situ*. Moreover, all animals that survived after administration of olaparib–sHDL–CpG nanoparticles did not develop tumors after being rechallenged in the contralateral hemisphere, indicating that the treatment elicited an immunological (adaptive) memory strong enough to eradicate the reimplanted tumor cells. Because all treatments are administered intracranially, differential systemic delivery is not a confounding factor. Instead, the benefit of the sHDL platform arises from coordinated payload release and immune stimulation. We show that CpG incorporation significantly enhances immune activation relative to olaparib alone, and prior work from our group has established the capacity of sHDL nanoparticles to improve payload bioactivity ^[23]^. The observed therapeutic benefit therefore demonstrates combined immune modulation and optimized payload presentation.

In conclusion, we provide preclinical evidence of the therapeutic efficacy of sHDL nanoparticles combining a PARP inhibitor (olaparib) and an immunostimulant agent (CpG oligodeoxynucleotides) for G34-mutant DHG. HDLs do not trespass the blood-brain barrier, thus we administered the formulation *in situ*. We propose that maximal safe surgery provides a window to administer the nanoparticle formulation *in situ* to patients. Surgery is usually followed by radiotherapy in DHG; thus this approach would maximize the therapeutic effect of the dual-drug nanoparticle formulation in combination with irradiation-induced DNA damage.

## Author contributions

SH, KB, AAM, TH, MS, SR performed experiments. SH and KB performed the western blot experiments. SH, KB, AAM, SS, PRL, AS and MGC analyzed the data. SH, KB, AAM, SS, PRL, AS and MGC designed the figures. SH, PRL, AS and MGC designed the research and wrote and edited the manuscript. All the authors revised the manuscript.

## Acknowledgments

This work was supported by the National Institutes of Health (NIH)/National Institute of Neurological Disorders and Stroke (NIH/NINDS) grants R01-NS1221165-01-A1; R01-NS122536; R01-NS124167 to MGC and PRL, and Rogel Cancer Center Scholar Award to M.G. Castro. This work has also been supported by NIH grants R01-HL165688 to A. Schwendeman, as well as the Department of Neurosurgery, the Pediatric Brain Tumor Foundation, The Pediatric Cancer Foundation, Leah’s Happy Hearts Foundation, Ian’s Friends Foundation (IFF), Chad Tough Foundation, and Smiles for Sophie Forever Foundation to MGC and PRL. We would also like to acknowledge the University of Michigan In-Vivo Animal Core (IVAC) facility and University of Michigan Institutional Animal Care and Use Committee (IACUC).

Address correspondence to: Maria G. Castro, University of Michigan Medical School, Department of Neurosurgery, 1150 West Medical Center Drive, MSRB II, Room 4570, Ann Arbor, Michigan 48109, USA. Phone: 734.764.0850.

## Data Availability Statement

The data that supports the findings of this study are available from the corresponding author upon reasonable request.

## RNA-Seq data availability

The RNA-Seq data sets have been deposited in the NCBI’s Gene Expression Omnibus (GEO) database (GEO GSE182068 and GSE182069).

## Phospho Antibody Array data availability

The full differential analysis data from the Phospho Antibody Arrays presented in this manuscript are available as supplementary tables (Tables 1 and 2)

## Graphical abstract

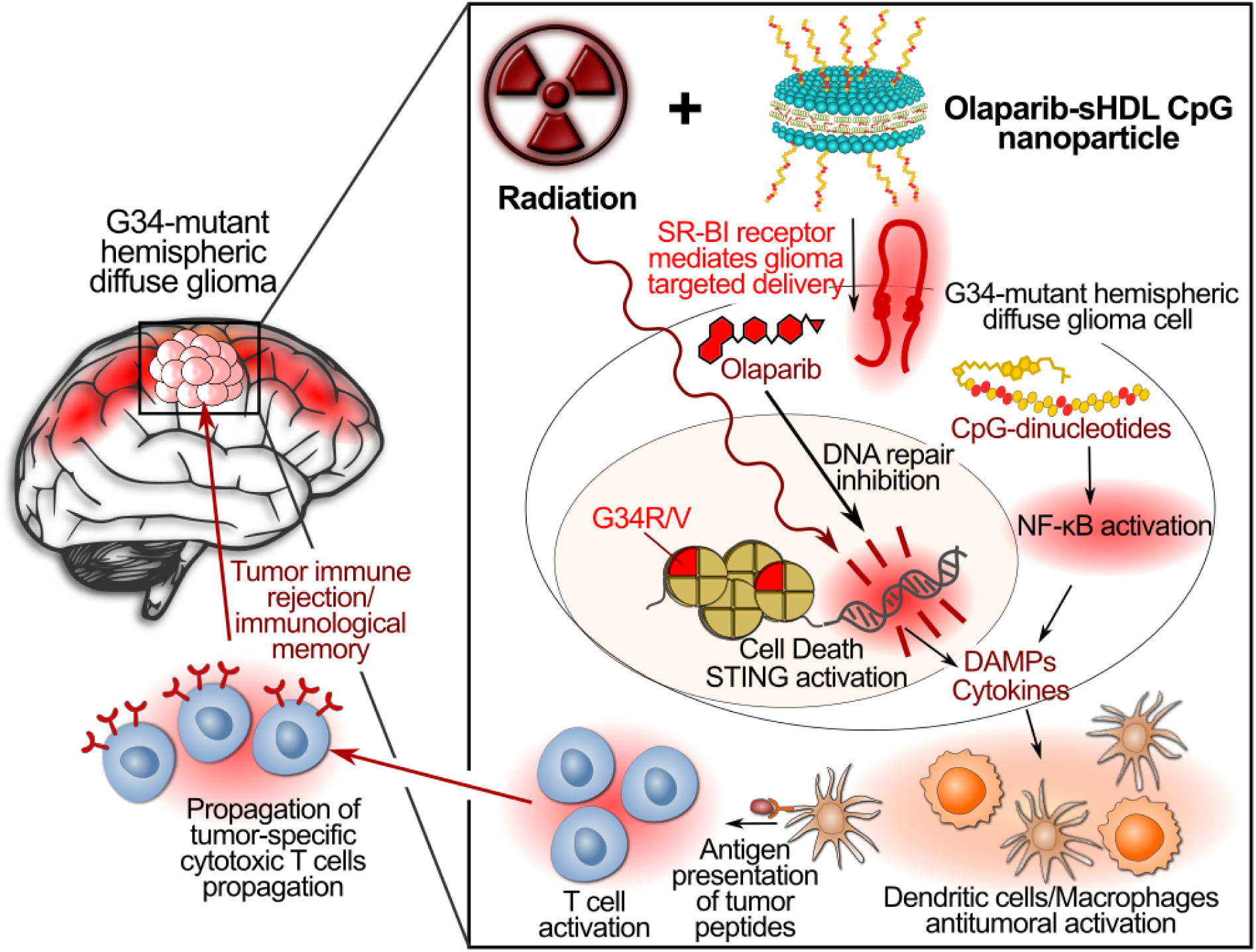

## Supplementary Figures

**Supplementary Figure 1.**
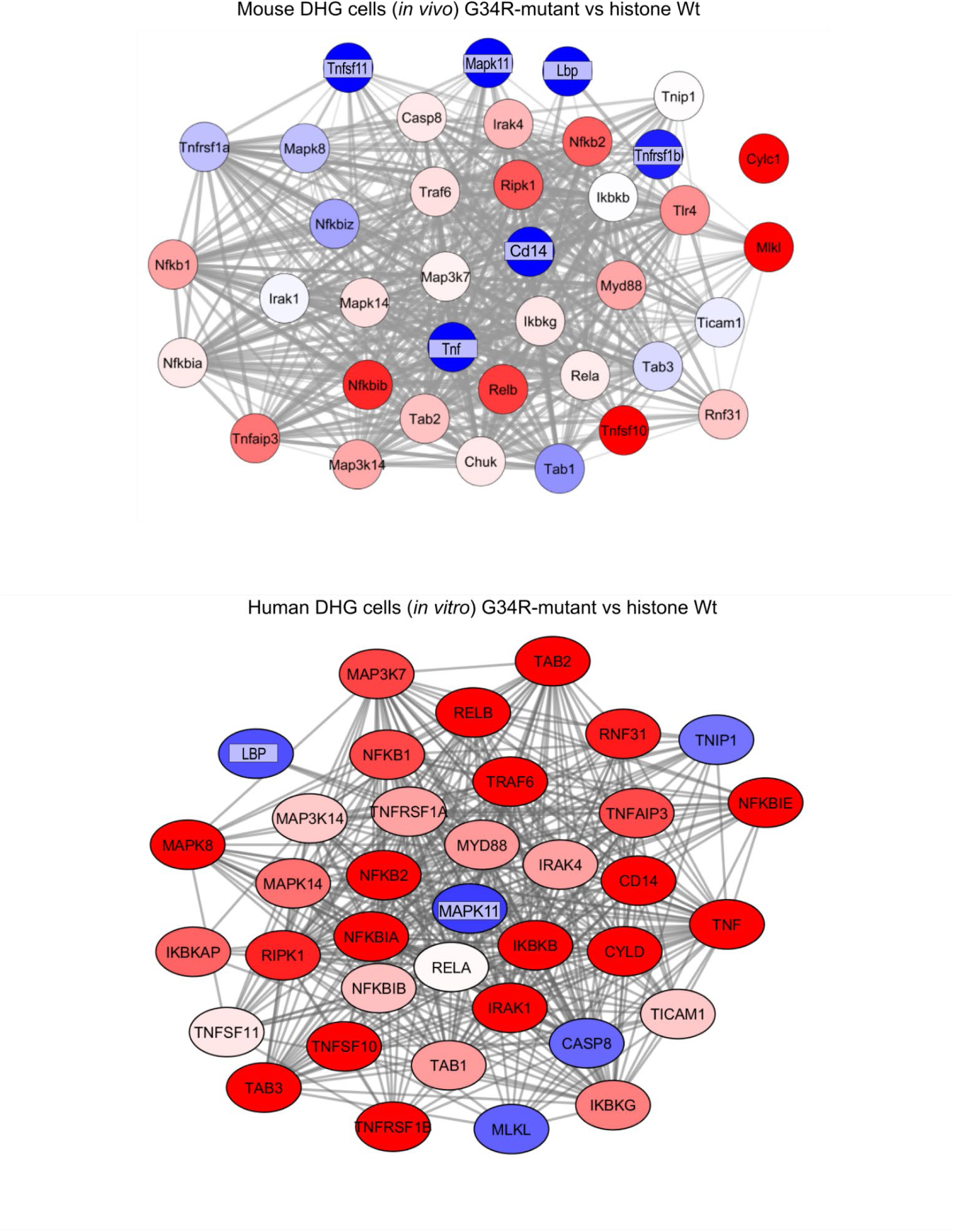
Transcriptional expression of NF-κB-related genes in G34R-mutant versus histone Wt mouse and human DHG cells.

**Supplementary figure 2.**
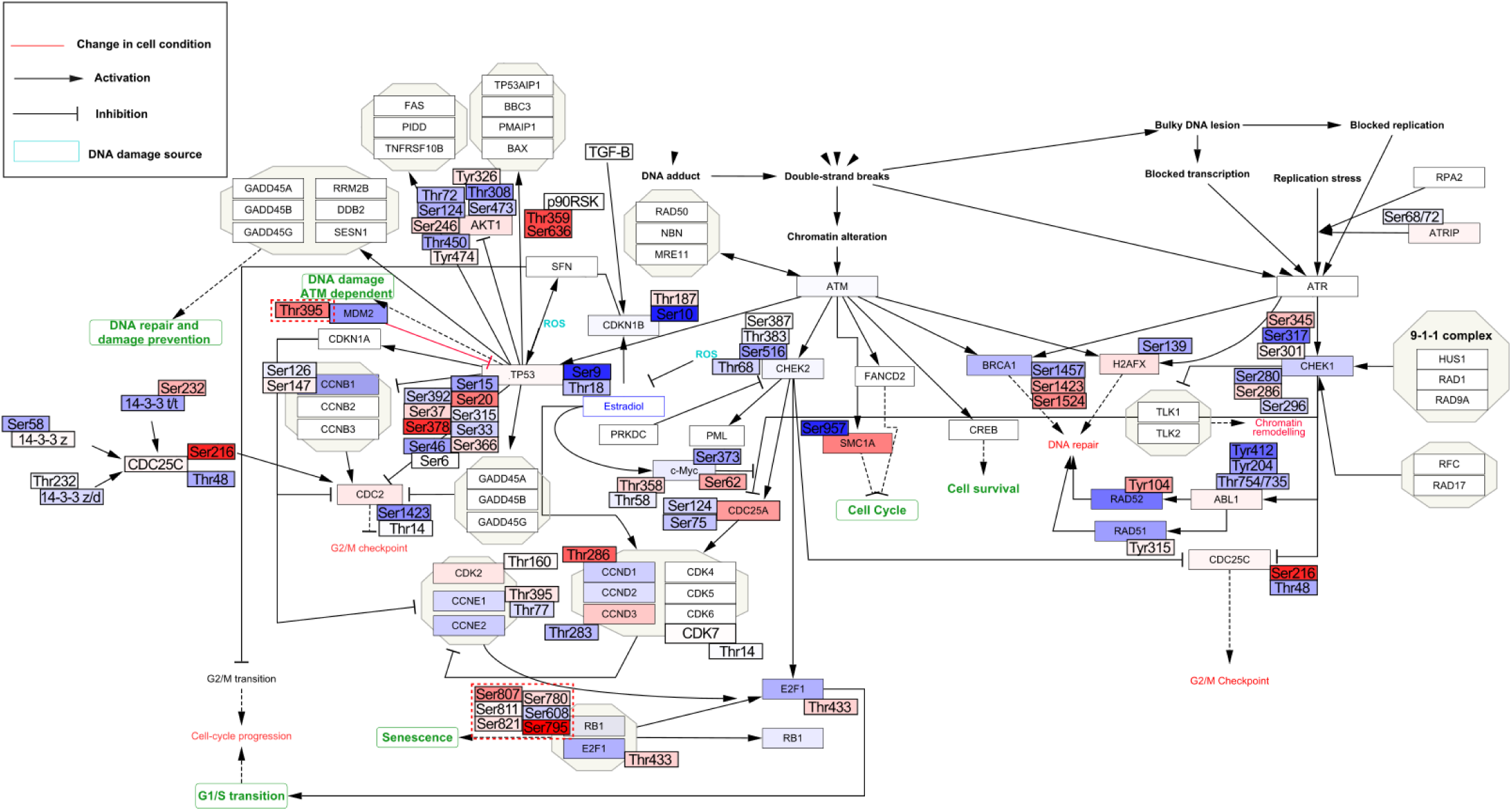
Postraslational activation of DNA repair and cell cycle genes in G34R mouse pHGG cells after treatment with olaparib-sHDL-CpG nanoparticles

**Supplementary Figure 3.**
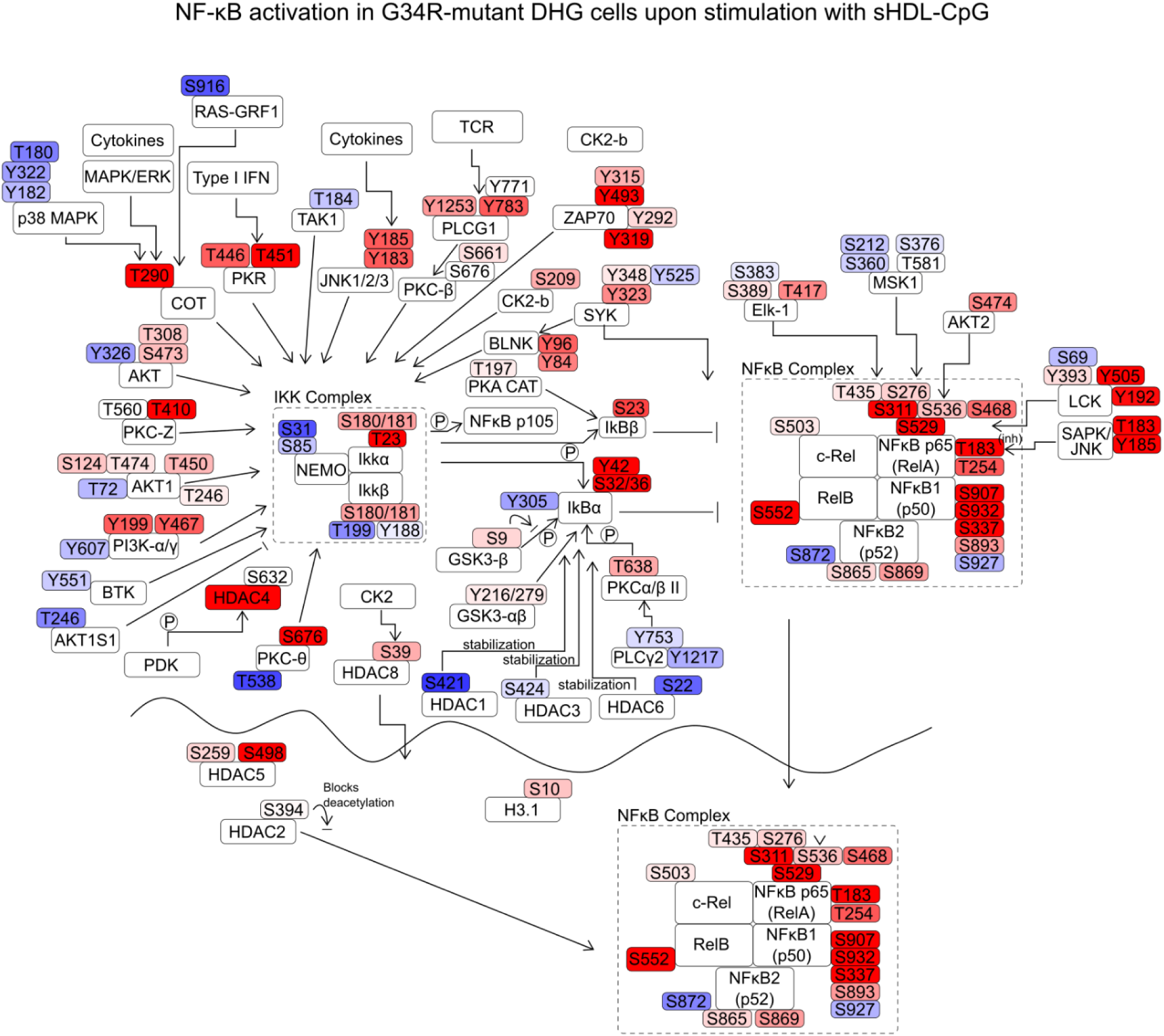
Postraslational activation of NF-kB genes in G34R-mutant DHG cells upon stimulation with sHDL-CpG nanoparticles

**Supplementary Figure 4.**
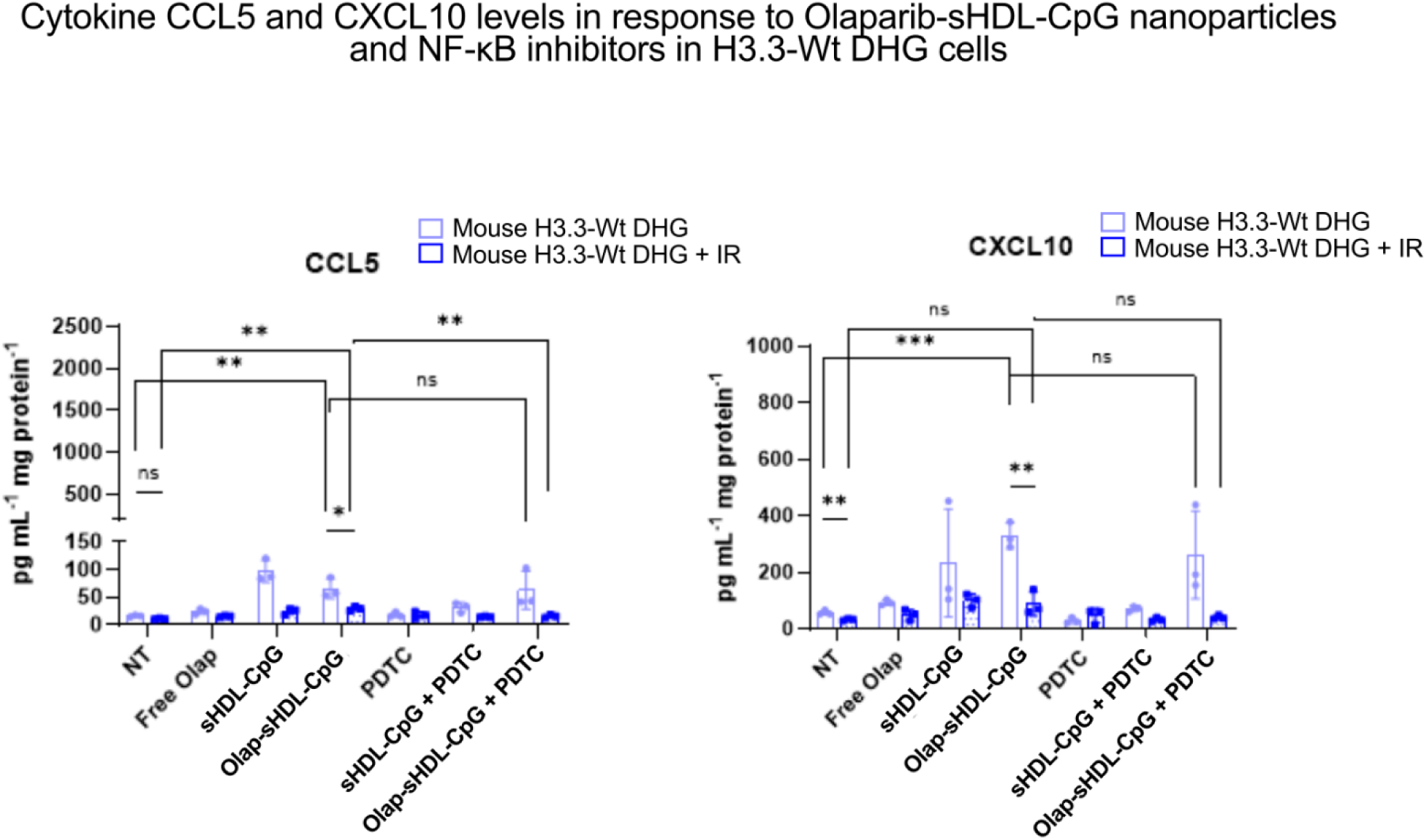
CCL5 and CXCL10 cytokine levels on H3.3-Wt mouse DHG cells upon stimulation with olaparib-sHDL-CpG nanoparticles in presence or absence of NF-κB inhibitors.

**Supplementary Figure 5.**
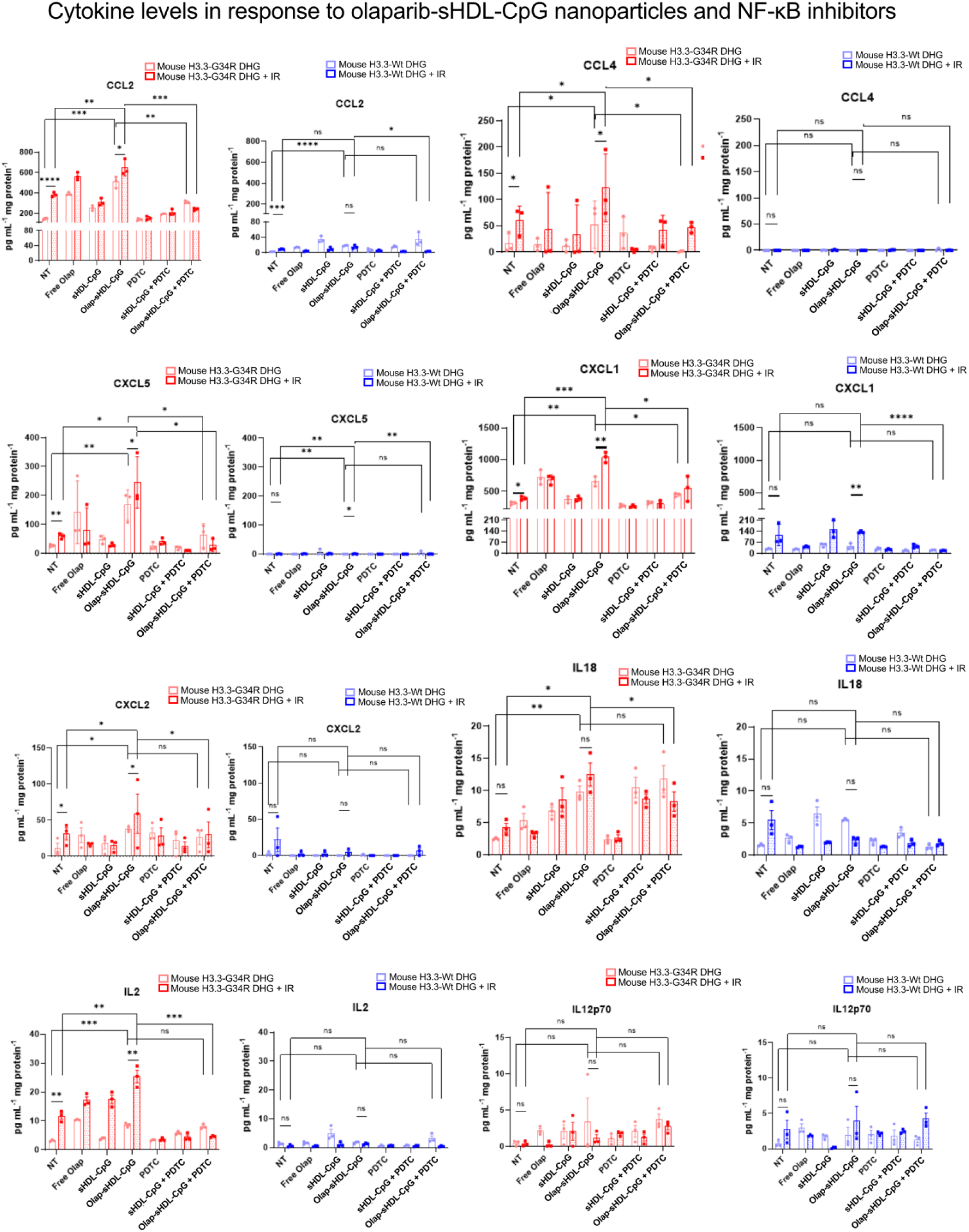
Cytokine levels on G34R-mutant and H3.3-Wt mouse DHG cells upon stimulation with olaparib-sHDL-CpG nanoparticles in presence or absence of NF-κB inhibitors.

**Supplementary Figure 6.**
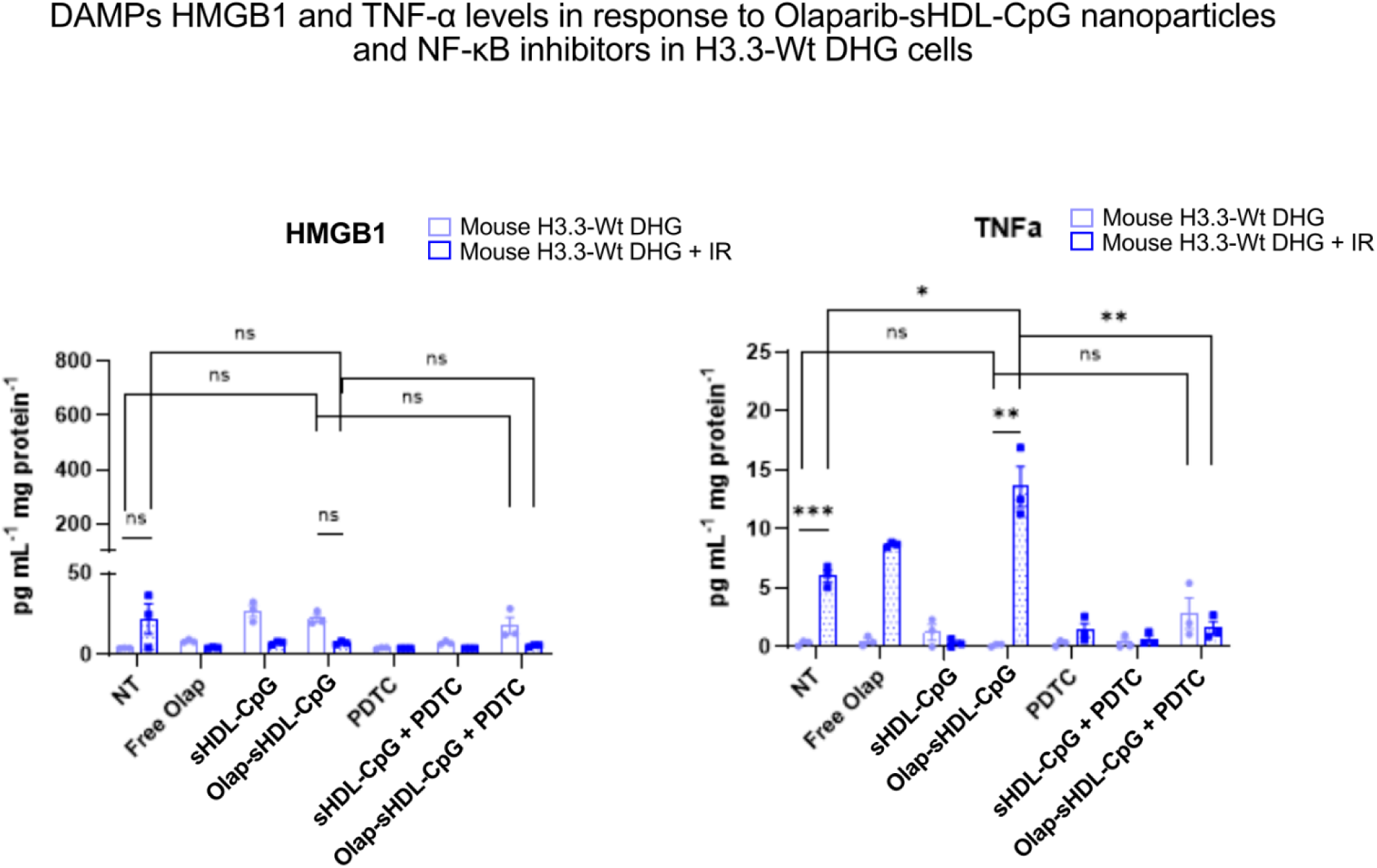
HMGB1 and TNFα DAMPs levels on H3.3-Wt mouse DHG cells upon stimulation with olaparib-sHDL-CpG nanoparticles in presence or absence of NF-κB inhibitors.

**Supplementary Figure 7.**
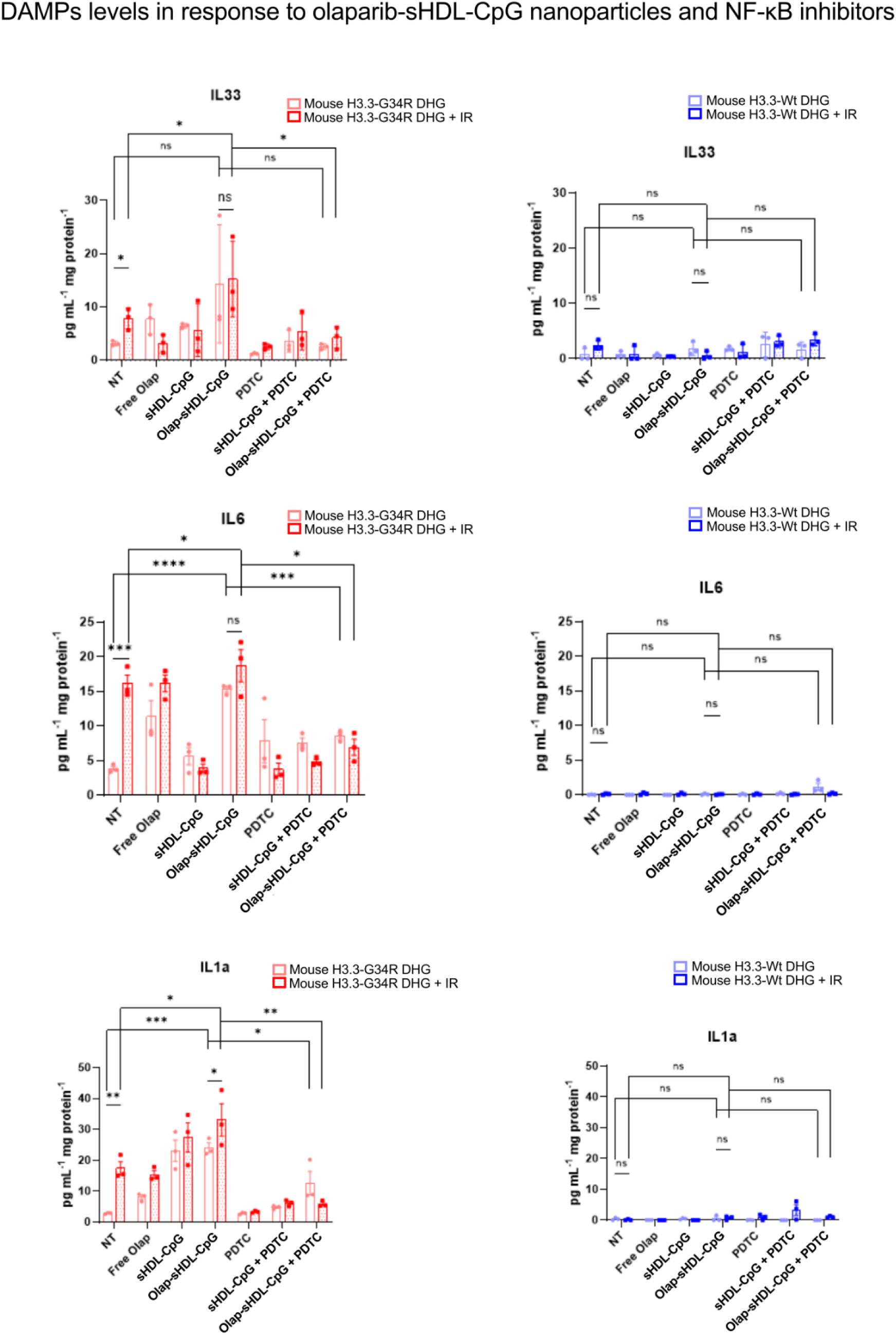
HMGB1 and TNFα DAMPs levels on G34R-mutant and H3.3-Wt mouse DHG cells upon stimulation with olaparib-sHDL-CpG nanoparticles in presence or absence of NF-κB inhibitors.

**Supplementary Figure 8.**
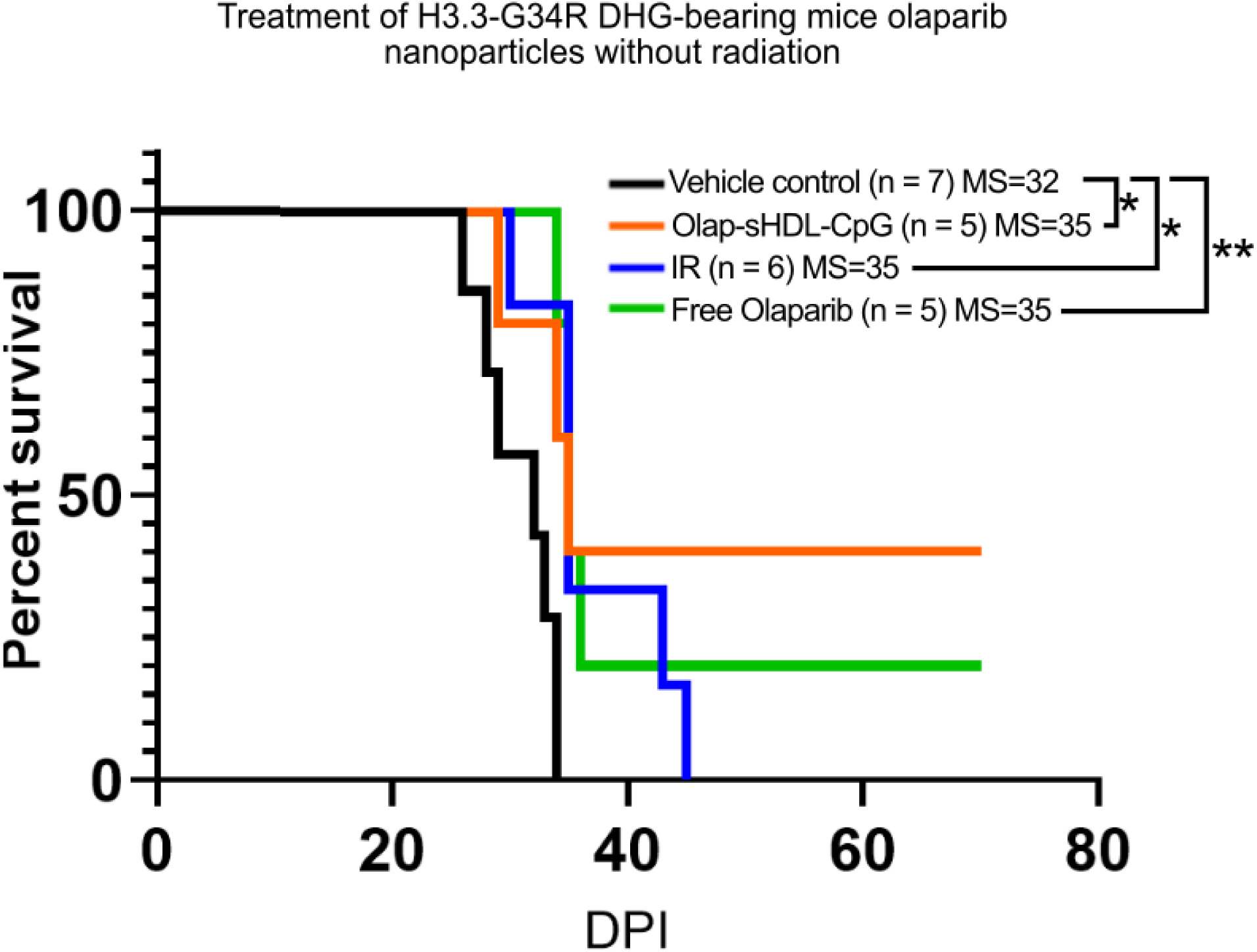
Survival curves H3.3-G34R DHG-bearing mice treated with olaparib-sHDL-CpG nanoparticles or free olaparib without radiation.

**Supplementary Figure 9.**
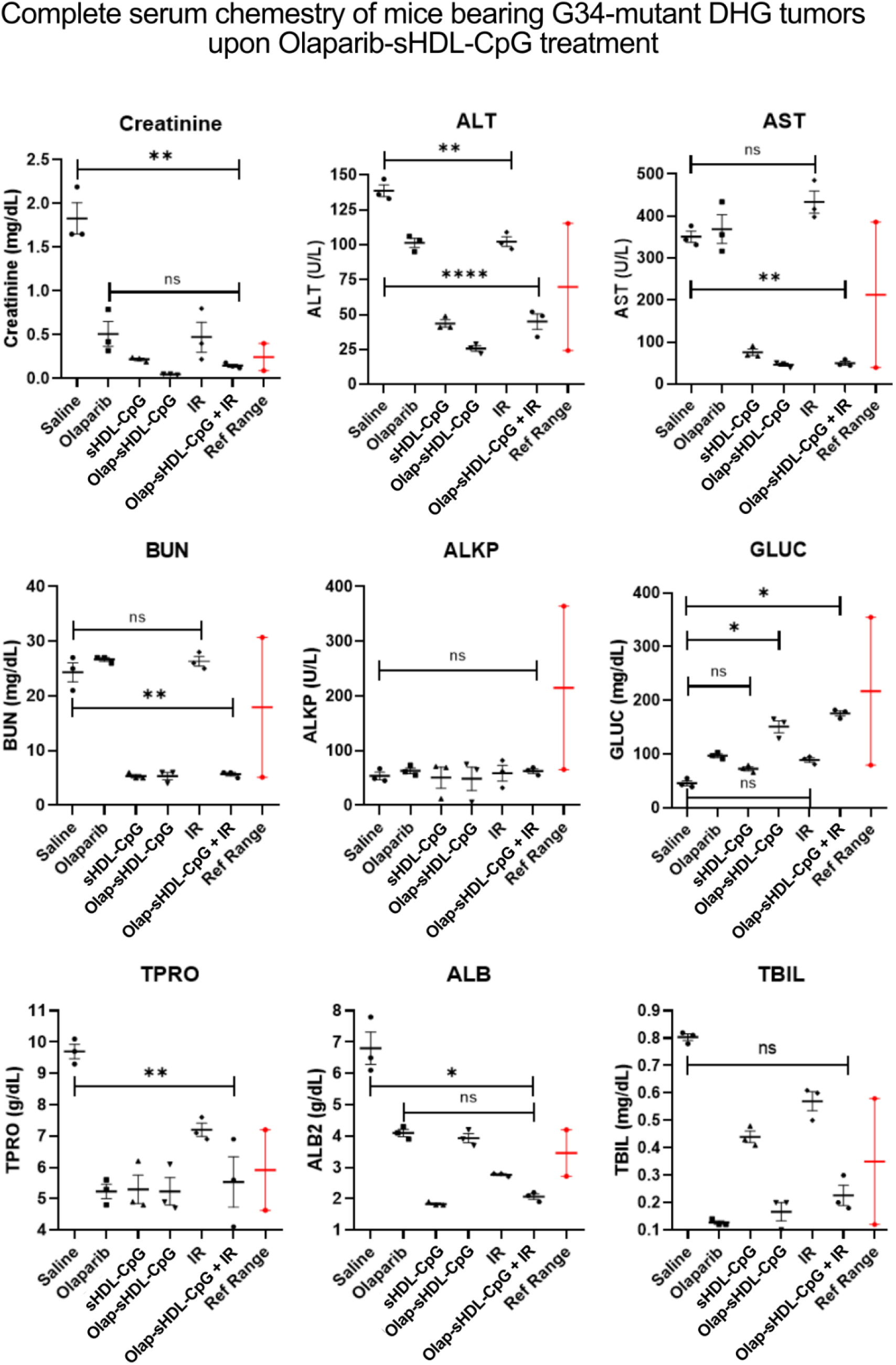
Assessment of liver function via evaluation of serum chemistry markers on H3.3-G34R DHG-bearing mice treated with olaparib-sHDL-CpG nanoparticles.

**Supplementary Figure 10.**
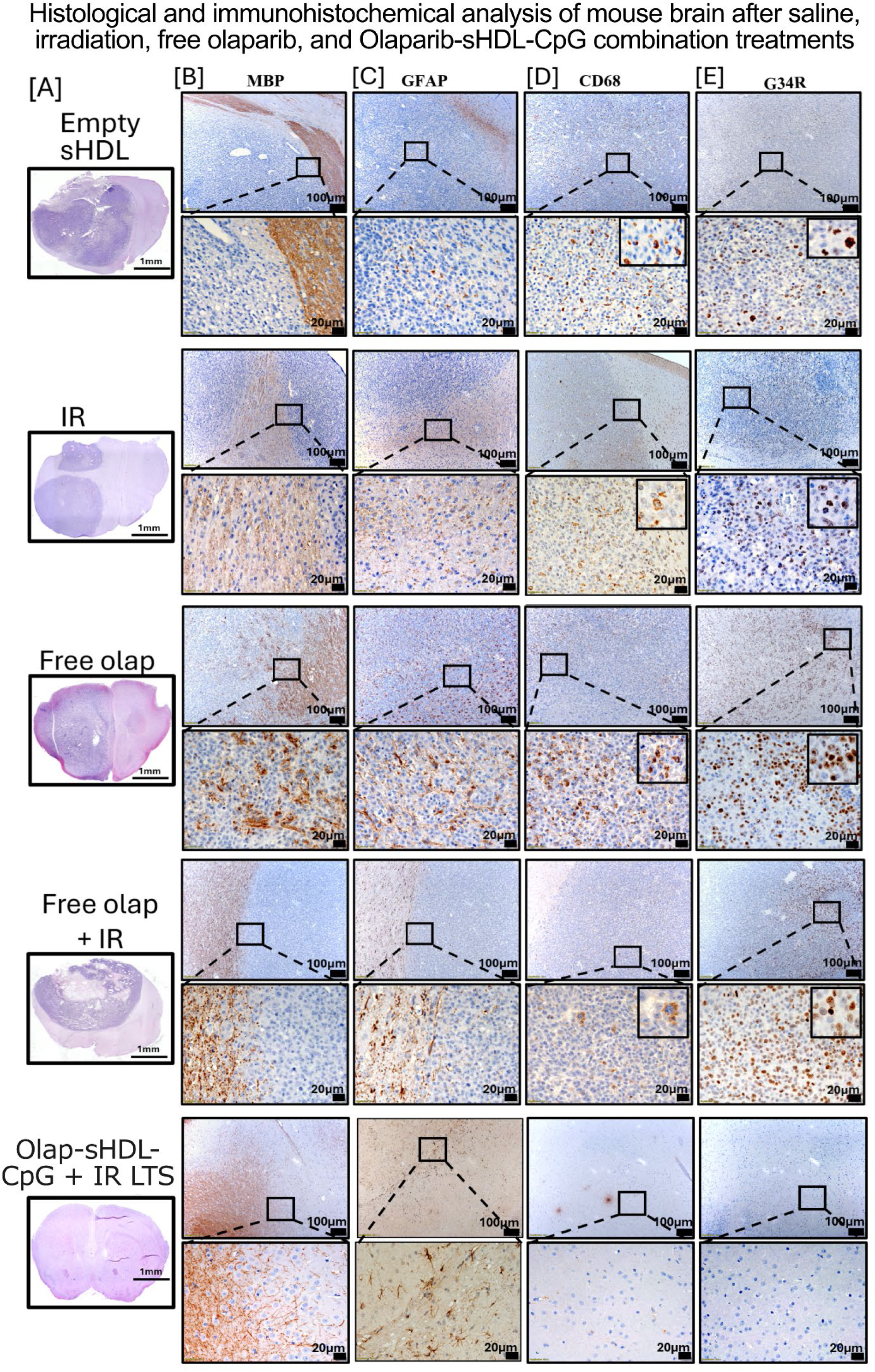
IHC tumor microenvironment and brain inflammation analysis on brains of H3.3-G34R DHG-bearing mice treated with olaparib-sHDL-CpG nanoparticles. Treatment group is indicated in (A) and antibody stains are indicated in (B) (MBP = Myelin basic protein), (C) (GFAP = glial fibrillary acidic protein), (D) (CD68 = CD68 glycoprotein, primarily expressed on macrophages and monocytes), and (E) (G34R= Anti H3.3 histone G34R-mutant)

**Supplementary Figure 11.**
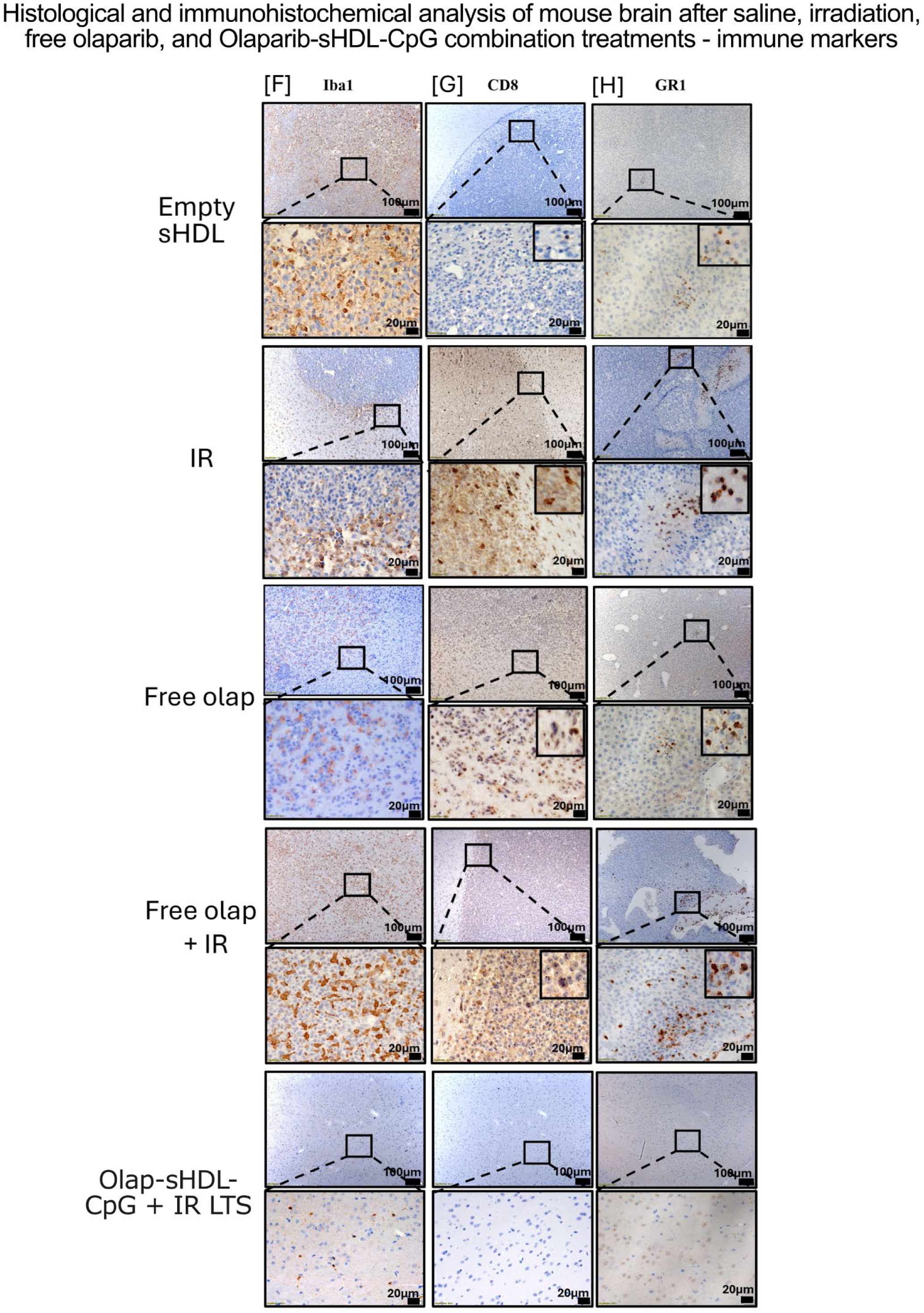
IHC tumor microenvironment and brain inflammation analysis on brains of H3.3-G34R DHG-bearing mice treated with olaparib-sHDL-CpG nanoparticles. Antibody stains are indicated in (A) (IBA1 = ionized calcium-binding adapter molecule 1, marker for microglia and macrophages), (B), (CD8 = CD8 antigen, a cell-surface glycoprotein primarily found on cytotoxic T-lymphocytes (CTLs)), and (C) (Gr1 = targets the Ly-6G antigen, a myeloid differentiation marker primarily found on neutrophils, monocytes, and granulocytes).

## References

[1] Q. T. Ostrom, H. Gittleman, G. Truitt, A. Boscia, C. Kruchko, J. S. Barnholtz-Sloan, Neuro Oncol. 2018, 20, iv1.

[2] C. Jones, M. A. Karajannis, D. T. W. Jones, M. W. Kieran, M. Monje, S. J. Baker, O. J. Becher, Y.-J. Cho, N. Gupta, C. Hawkins, D. Hargrave, D. A. Haas-Kogan, N. Jabado, X.-N. Li, S. Mueller, T. Nicolaides, R. J. Packer, A. I. Persson, J. J. Phillips, E. F. Simonds, J. M. Stafford, Y. Tang, S. M. Pfister, W. A. Weiss, Neuro Oncol. 2017, 19, 153.

[3] G. L. Lin, K. M. Wilson, M. Ceribelli, B. Z. Stanton, P. J. Woo, S. Kreimer, E. Y. Qin, X. Zhang, J. Lennon, S. Nagaraja, P. J. Morris, M. Quezada, S. M. Gillespie, D. Y. Duveau, A. M. Michalowski, P. Shinn, R. Guha, M. Ferrer, C. Klumpp-Thomas, S. Michael, C. McKnight, P. Minhas, Z. Itkin, E. H. Raabe, L. Chen, R. Ghanem, A. C. Geraghty, L. Ni, K. I. Andreasson, N. A. Vitanza, K. E. Warren, C. J. Thomas, M. Monje, Sci. Transl. Med. 2019, 11, DOI 10.1126/scitranslmed.aaw0064.

[4] Z. Miklja, A. Pasternak, S. Stallard, T. Nicolaides, C. Kline-Nunnally, B. Cole, R. Beroukhim, P. Bandopadhayay, S. Chi, S. H. Ramkissoon, B. Mullan, A. K. Bruzek, A. Gauthier, T. Garcia, C. Atchison, B. Marini, M. Fouladi, D. W. Parsons, S. Leary, S. Mueller, K. L. Ligon, C. Koschmann, Neuro Oncol. 2019, 21, 968.

[5] R. L. Siegel, K. D. Miller, N. S. Wagle, A. Jemal, CA Cancer J. Clin. 2023, 73, 17.

[6] A. Mackay, A. Burford, D. Carvalho, E. Izquierdo, J. Fazal-Salom, K. R. Taylor, L. Bjerke, M. Clarke, M. Vinci, M. Nandhabalan, S. Temelso, S. Popov, V. Molinari, P. Raman, A. J. Waanders, H. J. Han, S. Gupta, L. Marshall, S. Zacharoulis, S. Vaidya, H. C. Mandeville, L. R. Bridges, A. J. Martin, S. Al-Sarraj, C. Chandler, H.-K. Ng, X. Li, K. Mu, S. Trabelsi, D. H.-B. Brahim, A. N. Kisljakov, D. M. Konovalov, A. S. Moore, A. M. Carcaboso, M. Sunol, C. de Torres, O. Cruz, J. Mora, L. I. Shats, J. N. Stavale, L. T. Bidinotto, R. M. Reis, N. Entz-Werle, M. Farrell, J. Cryan, D. Crimmins, J. Caird, J. Pears, M. Monje, M.-A. Debily, D. Castel, J. Grill, C. Hawkins, H. Nikbakht, N. Jabado, S. J. Baker, S. M. Pfister, D. T. W. Jones, M. Fouladi, A. O. von Bueren, M. Baudis, A. Resnick, C. Jones, Cancer Cell 2017, 32, 520.

[7] J. Schwartzentruber, A. Korshunov, X.-Y. Liu, D. T. W. Jones, E. Pfaff, K. Jacob, D. Sturm, A. M. Fontebasso, D.-A. K. Quang, M. Tönjes, V. Hovestadt, S. Albrecht, M. Kool, A. Nantel, C. Konermann, A. Lindroth, N. Jäger, T. Rausch, M. Ryzhova, J. O. Korbel, T. Hielscher, P. Hauser, M. Garami, A. Klekner, L. Bognar, M. Ebinger, M. U. Schuhmann, W. Scheurlen, A. Pekrun, M. C. Frühwald, W. Roggendorf, C. Kramm, M. Dürken, J. Atkinson, P. Lepage, A. Montpetit, M. Zakrzewska, K. Zakrzewski, P. P. Liberski, Z. Dong, P. Siegel, A. E. Kulozik, M. Zapatka, A. Guha, D. Malkin, J. Felsberg, G. Reifenberger, A. von Deimling, K. Ichimura, V. P. Collins, H. Witt, T. Milde, O. Witt, C. Zhang, P. Castelo-Branco, P. Lichter, D. Faury, U. Tabori, C. Plass, J. Majewski, S. M. Pfister, N. Jabado, Nature 2012, 482, 226.

[8] D. Sturm, H. Witt, V. Hovestadt, D.-A. Khuong-Quang, D. T. W. Jones, C. Konermann, E. Pfaff, M. Tönjes, M. Sill, S. Bender, M. Kool, M. Zapatka, N. Becker, M. Zucknick, T. Hielscher, X.-Y. Liu, A. M. Fontebasso, M. Ryzhova, S. Albrecht, K. Jacob, M. Wolter, M. Ebinger, M. U. Schuhmann, T. van Meter, M. C. Frühwald, H. Hauch, A. Pekrun, B. Radlwimmer, T. Niehues, G. von Komorowski, M. Dürken, A. E. Kulozik, J. Madden, A. Donson, N. K. Foreman, R. Drissi, M. Fouladi, W. Scheurlen, A. von Deimling, C. Monoranu, W. Roggendorf, C. Herold-Mende, A. Unterberg, C. M. Kramm, J. Felsberg, C. Hartmann, B. Wiestler, W. Wick, T. Milde, O. Witt, A. M. Lindroth, J. Schwartzentruber, D. Faury, A. Fleming, M. Zakrzewska, P. P. Liberski, K. Zakrzewski, P. Hauser, M. Garami, A. Klekner, L. Bognar, S. Morrissy, F. Cavalli, M. D. Taylor, P. van Sluis, J. Koster, R. Versteeg, R. Volckmann, T. Mikkelsen, K. Aldape, G. Reifenberger, V. P. Collins, J. Majewski, A. Korshunov, P. Lichter, C. Plass, N. Jabado, S. M. Pfister, Cancer Cell 2012, 22, 425.

[9] L. Bjerke, A. Mackay, M. Nandhabalan, A. Burford, A. Jury, S. Popov, D. A. Bax, D. Carvalho, K. R. Taylor, M. Vinci, I. Bajrami, I. M. McGonnell, C. J. Lord, R. M. Reis, D. Hargrave, A. Ashworth, P. Workman, C. Jones, Cancer Discov. 2013, 3, 512.

[10] J. Puntonet, V. Dangouloff-Ros, R. Saffroy, M. Pagès, F. Andreiuolo, J. Grill, S. Puget, N. Boddaert, P. Varlet, J. Neuroradiol. 2018, 45, 316.

[11] K. Yoshimoto, R. Hatae, Y. Sangatsuda, S. O. Suzuki, N. Hata, Y. Akagi, D. Kuga, M. Hideki, K. Yamashita, O. Togao, A. Hiwatashi, T. Iwaki, M. Mizoguchi, K. Iihara, Brain Tumor Pathol. 2017, 34, 103.

[12] P. Pathak, P. Jha, S. Purkait, V. Sharma, V. Suri, M. C. Sharma, M. Faruq, A. Suri, C. Sarkar, J Neurooncol 2015, 121, 489.

[13] G. Wu, A. Broniscer, T. A. McEachron, C. Lu, B. S. Paugh, J. Becksfort, C. Qu, L. Ding, R. Huether, M. Parker, J. Zhang, A. Gajjar, M. A. Dyer, C. G. Mullighan, R. J. Gilbertson, E. R. Mardis, R. K. Wilson, J. R. Downing, D. W. Ellison, J. Zhang, S. J. Baker, St. Jude Children’s Research Hospital–Washington University Pediatric Cancer Genome Project, Nat. Genet. 2012, 44, 251.

[14] A. Korshunov, D. Capper, D. Reuss, D. Schrimpf, M. Ryzhova, V. Hovestadt, D. Sturm, J. Meyer, C. Jones, O. Zheludkova, E. Kumirova, A. Golanov, M. Kool, U. Schüller, M. Mittelbronn, M. Hasselblatt, J. Schittenhelm, G. Reifenberger, C. Herold-Mende, P. Lichter, A. von Deimling, S. M. Pfister, D. T. W. Jones, Acta Neuropathol. 2016, 131, 137.

[15] A. Mackay, A. Burford, V. Molinari, D. T. W. Jones, E. Izquierdo, J. Brouwer-Visser, F. Giangaspero, C. Haberler, T. Pietsch, T. S. Jacques, D. Figarella-Branger, D. Rodriguez, P. S. Morgan, P. Raman, A. J. Waanders, A. C. Resnick, M. Massimino, M. L. Garrè, H. Smith, D. Capper, S. M. Pfister, T. Würdinger, R. Tam, J. Garcia, M. D. Thakur, G. Vassal, J. Grill, T. Jaspan, P. Varlet, C. Jones, Cancer Cell 2018, 33, 829.

[16] S. Haase, K. Banerjee, A. A. Mujeeb, C. S. Hartlage, F. M. Núñez, F. J. Núñez, M. S. Alghamri, P. Kadiyala, S. Carney, M. N. Barissi, A. W. Taher, E. K. Brumley, S. Thompson, J. T. Dreyer, C. T. Alindogan, M. B. Garcia-Fabiani, A. Comba, S. Venneti, V. Ravikumar, C. Koschmann, Á. M. Carcaboso, M. Vinci, A. Rao, J. S. Yu, P. R. Lowenstein, M. G. Castro, J. Clin. Invest. 2022, 132, DOI 10.1172/JCI154229.

[17] B. L. McClellan, S. Haase, F. J. Nunez, M. S. Alghamri, A. A. Dabaja, P. R. Lowenstein, M. G. Castro, J. Clin. Invest. 2023, 133, DOI 10.1172/JCI163450.

[18] N. Kamran, P. Kadiyala, M. Saxena, M. Candolfi, Y. Li, M. A. Moreno-Ayala, N. Raja, D. Shah, P. R. Lowenstein, M. G. Castro, Mol. Ther. 2017, 25, 232.

[19] M. S. Alghamri, B. L. McClellan, C. S. Hartlage, S. Haase, S. M. Faisal, R. Thalla, A. Dabaja, K. Banerjee, S. V. Carney, A. A. Mujeeb, M. R. Olin, J. J. Moon, A. Schwendeman, P. R. Lowenstein, M. G. Castro, Front. Pharmacol. 2021, 12, 680021.

[20] M. B. Garcia-Fabiani, S. Haase, K. Banerjee, Z. Zhu, B. L. McClellan, A. A. Mujeeb, Y. Li, C. E. Tronrud, M. L. Varela, M. E. J. West, J. Yu, P. Kadiyala, A. W. Taher, F. J. Núñez, M. S. Alghamri, A. Comba, F. M. Mendez, A. J. Nicola Candia, B. Salazar, F. M. Nunez, M. B. Edwards, T. Qin, R. T. Cartaxo, M. Niculcea, C. Koschmann, S. Venneti, M. P. Vallcorba, E. Nasajpour, G. Pericoli, M. Vinci, C. L. Kleinman, N. Jabado, J. P. Chandler, A. M. Sonabend, M. DeCuypere, D. Hambardzumyan, L. M. Prolo, K. B. Mahaney, G. A. Grant, C. K. Petritsch, J. D. Welch, M. A. Sartor, P. R. Lowenstein, M. G. Castro, BioRxiv 2025, DOI 10.1101/2023.06.13.544658.

[21] C. C. L. Chen, S. Deshmukh, S. Jessa, D. Hadjadj, V. Lisi, A. F. Andrade, D. Faury, W. Jawhar, R. Dali, H. Suzuki, M. Pathania, D. A, F. Dubois, E. Woodward, S. Hébert, M. Coutelier, J. Karamchandani, S. Albrecht, S. Brandner, N. De Jay, T. Gayden, A. Bajic, A. S. Harutyunyan, D. M. Marchione, L. G. Mikael, N. Juretic, M. Zeinieh, C. Russo, N. Maestro, A. V. Bassenden, P. Hauser, J. Virga, L. Bognar, A. Klekner, M. Zapotocky, A. Vicha, L. Krskova, K. Vanova, J. Zamecnik, D. Sumerauer, P. G. Ekert, D. S. Ziegler, B. Ellezam, M. G. Filbin, M. Blanchette, J. R. Hansford, D.-A. Khuong-Quang, A. M. Berghuis, A. G. Weil, B. A. Garcia, L. Garzia, S. C. Mack, R. Beroukhim, K. L. Ligon, M. D. Taylor, P. Bandopadhayay, C. Kramm, S. M. Pfister, A. Korshunov, D. Sturm, D. T. W. Jones, P. Salomoni, C. L. Kleinman, N. Jabado, Cell 2020, 183, 1617.

[22] S. Khazaei, C. C. L. Chen, A. F. Andrade, N. Kabir, P. Azarafshar, S. M. Morcos, J. A. França, M. Lopes, P. J. Lund, G. Danieau, S. Worme, L. Adnani, N. Nzirorera, X. Chen, G. Yogarajah, C. Russo, M. Zeinieh, C. J. Wong, L. Bryant, S. Hébert, B. Tong, T. S. Sihota, D. Faury, E. Puligandla, W. Jawhar, V. Sandy, M. Cowan, E. M. Nakada, L. A. Jerome-Majewska, B. Ellezam, C. C. Gomes, J. Denecke, D. Lessel, M. T. McDonald, C. E. Pizoli, K. Taylor, B. T. Cocanougher, E. J. Bhoj, A.-C. Gingras, B. A. Garcia, C. Lu, E. I. Campos, C. L. Kleinman, L. Garzia, N. Jabado, Cell 2023, 186, 1162.

[23] P. Kadiyala, D. Li, F. M. Nuñez, D. Altshuler, R. Doherty, R. Kuai, M. Yu, N. Kamran, M. Edwards, J. J. Moon, P. R. Lowenstein, M. G. Castro, A. Schwendeman, ACS Nano 2019, 13, 1365.

[24] C. P. Hall, J. C. Cronk, J. A. Rubens, J. Clin. Invest. 2022, 132, DOI 10.1172/JCI164420.

[25] W.-J. Shen, J. Hu, Z. Hu, F. B. Kraemer, S. Azhar, Metab. Clin. Exp. 2014, 63, 875.

[26] R. L. Bowman, Q. Wang, A. Carro, R. G. W. Verhaak, M. Squatrito, Neuro Oncol. 2017, 19, 139.

[27] T. Liu, L. Zhang, D. Joo, S.-C. Sun, Signal Transduct. Target. Ther. 2017, 2, 17023.

[28] C. A. Schneider, W. S. Rasband, K. W. Eliceiri, Nat. Methods 2012, 9, 671.

[29] G. Carpentier, E. Henault, in Proceedings of the ImageJ User and Developer Conference, Centre de Recherche Public Henri Tudor, Ed., 2010, p. 238–240.

[30] M. A. Karajannis, A. Onar-Thomas, T. Lin, P. A. Baxter, D. R. Boué, B. L. Cole, C. Fuller, S. Haque, N. Jabado, J. T. Lucas, S. M. MacDonald, C. Matsushima, N. Patel, C. R. Pierson, M. M. Souweidane, D. L. Thomas, M. F. Walsh, W. Zaky, S. E. S. Leary, A. Gajjar, M. Fouladi, K. J. Cohen, Neuro Oncol. 2025, 27, 1092.

