## Supplemental Table 1 for "Targeting NF-κB epigenetic activation and DNA repair deficiency in G34-mutant pediatric diffuse hemispheric glioma with nanoparticles combining PARP inhibition and immune stimulation mediated by CpG dinucleotides"

| **Company (Cat#)** | **Primary Ab** | **Host** | **Dilutions** | **Secondary Ab** |
| --- | --- | --- | --- | --- |
| Millipore (MAB386) | Anti-MBP  (a.a. 82-87) | Rat | 1:500 | Anti-Rat Biotinylated  (1:1000) |
| Millipore  (AB5541) | Anti-GFAP | Chicken | 1:1000 | Anti-Chicken  Biotinylated  (1:1000) |
| Abcam (ab125212) | Anti-CD68 | Rabbit | 1:200 | Anti-Rabbit Biotinylated  (1:1000) |
| Revmab biosciences  31-1120-00-S | Anti-H3.3-G34R | Rabbit | 1:500 | Anti-Rabbit Biotinylated  (1:1000) |
| Abcam  (ab178846) | Anti-Iba1 | Rabbit | 1:2000 | Anti-Rabbit Biotinylated  (1:1000) |
| HistoSure  (HS-361 003) | Anti-CD8a  (230-247) | Rabbit | 1:100 | Anti-Rabbit Biotinylated  (1:1000) |
| InvitrogenMA5-32001 | SR-BI Antibody (SR37-06) | Rabbit | 1;100 | Anti-Rabbit Biotinylated  (1:1000) |
| Novus Biologicals  NB600-1387 | Anti-GR1 | Rat | 1:50 | Anti-Rat Biotinylated |
| Abcam | Anti-Scavenging Receptor SR-BI antibody [EP1556Y] | Rabbit monoclonal | 1:1000 | Anti-Rabbit HRP  (1:2000) |
| ThermoFisher Scientific | Phospho-NFkB p65 (Ser529) Polyclonal Antibody | Rabbit | 1:1000 | Anti-Rabbit HRP  (1:2000) |
| Cell Signaling Technology | NF-kappaB p65 (D14E12) Rabbit Monoclonal Antibody | Rabbit | 1:1000 | Anti-Rabbit HRP  (1:2000) |
| ThermoFisher Scientific | Phospho-NFkB p105/p50 (Ser907) Polyclonal Antibody | Rabbit | 1:1000 | Anti-Rabbit HRP  (1:2000) |
| Santa Cruz Biotechnology | NFκB p50 Antibody (E-10) | mouse monoclonal | 1:1000 | Anti-Mouse HRP  (1:2000) |
| Cell Signaling Technology | beta-Actin (8H10D10) Mouse Monoclonal Antibody | mouse monoclonal | 1:1000 | Anti-Mouse HRP  (1:2000) |
| Cell Signaling Technology | Phospho-IKK alpha/beta (Ser176/180) (16A6) Rabbit Monoclonal Antibody | Rabbit | 1:1000 | Anti-Rabbit HRP  (1:2000) |
| IKK beta (D30C6) Rabbit Monoclonal Antibody | IKK beta (D30C6) Rabbit Monoclonal Antibody | Rabbit | 1:1000 | Anti-Rabbit HRP  (1:2000) |

Supplementary Table 1 – List of antibodies
